# A rapidly closing window for coral persistence under global warming

**DOI:** 10.1101/2025.01.23.634487

**Authors:** Yves-Marie Bozec, Arne A.S. Adam, Beatriz Arellano Nava, Anna K. Cresswell, Vanessa Haller-Bull, Takuya Iwanaga, Liam Lachs, Samuel A. Matthews, Jennifer K. McWhorter, Kenneth R.N. Anthony, Scott A. Condie, Paul R. Halloran, Juan-Carlos Ortiz, Cynthia Riginos, Peter J. Mumby

**Affiliations:** School of the Environment, University of Queensland, St Lucia, Queensland, Australia; Global Systems Institute and Geography Department, University of Exeter, Exeter, UK; Australian Institute of Marine Science, Townsville, Queensland, Australia; Oceans Institute, University of Western Australia, Perth, Western Australia, Australia; School of Natural and Environmental Sciences, Newcastle University, Newcastle upon Tyne, UK; NOAA/OAR/Atlantic Oceanographic & Meteorological Laboratory, Miami, FL, 33149 USA; CSIRO Environment, Hobart, Tasmania, Australia

## Abstract

Coral reefs around the world are threatened by recurrent marine heatwaves causing mass coral bleaching and mortality^1^. Mitigating future warming impacts requires strategic management^2^ that adopts a long-term lens. Global analyses of projected heatwaves^3,4^ are critical for decision-making but how disturbance refugia, coral life-histories and adaptation interact with warming is unknown. Here, we simulate coral eco-evolutionary dynamics across >3,800 individual reefs of Australia’s iconic Great Barrier Reef under the current suite of climate projections^5^. We forecast a rapid coral decline before mid-century regardless of emissions scenario. Under the most likely warming^6^, corals may continue to decline throughout the century. However, corals may recover after mid-century provided that warming is sufficiently slow, allowing thermal adaptation to keep pace with temperature changes – as projected if warming does not exceed 2°C. Higher emission scenarios would drive most reefs to a near collapse. Future resilient reefs were mostly found in bleaching refugia, which also possessed a greater diversity of thermal phenotypes. Although cool-adapted corals disperse to thermal hotspots, there was no evidence of ‘gene swamping’ undermining thermal adaptation. While opportunities exist to build management strategies that promote adaptation in thermal refugia and warm spots, curbing the rate of warming by 2050 is crucial for coral persistence.

## Main text

Anthropogenic greenhouse-gas (GHG) emissions have raised global temperatures by roughly 1°C above pre-industrial levels^5^, intensifying the frequency and severity of marine heatwaves and driving tropical coral reef ecosystems into decline^7–9^. Heat stress disrupts the symbiotic relationship between reef-building corals and their photosynthetic algal endosymbionts (zooxanthellae), leading to coral bleaching^10^. Under severe and prolonged heat stress, persistent coral bleaching leads to extensive mortality^8,9,11^. Regional-scale marine heatwaves are occurring at an alarming rate, significantly reducing the recovery time between consecutive bleaching events^1,12,13^. This poses an unprecedented threat to the persistence of functioning coral reef ecosystems in the 21^st^ century: even with the most aggressive reduction of GHG emissions, global warming is expected to exceed 1.5°C for multiple decades^14^.

Projections of reef futures have focused almost entirely on evaluating the likelihood that temperatures will exceed current thresholds for widespread coral mortality^15,16^. These projections fail to account for evolutionary processes and the adaptive capacity of corals, the diversity of coral life-histories, and the complex demographics of interconnected coral metacommunities within a spatially heterogeneous environment, including disturbance refugia^16,17^. The ability of corals to withstand rising sea temperatures will partly depend on their adaptive potential (i.e., *evolutionary rescue*^18^), yet the degree to which natural selection can enhance heat tolerance under frequent and intense marine heatwaves is unknown. Population persistence might also be promoted by coral larvae supplied by nearby reef habitats (i.e., *demographic rescue*^19^), but the resilience of larval dispersal networks remains uncertain given the large geographic footprint of marine heatwaves^20^. The fate of coral reefs will ultimately hinge on how effectively these rescue effects may combine to enhance resistance to bleaching and support recovery, which in turn will be influenced heavily by the rate of temperature change in the next decades. Here, we explore the eco-evolutionary response of coral communities to climate change using a comprehensive and field-tested ecosystem model of Australia’s Great Barrier Reef^21^ extended to include adaptive evolution to thermal stress. By simulating community evolutionary dynamics throughout the 21^st^ century, we evaluate coral persistence under alternative scenarios of GHG emissions and assess the potential for rapid coral evolution across multiple spatial and temporal scales. While our findings paint a grim future for coral reefs, we identify paths under which coral populations may adapt and persist. We also find opportunities for management interventions to capitalise on coral connectivity patterns and anticipated future warming (thermal refugia and warm spots) for supporting reef resilience, reinforcing the value of taking a strategic and proactive approach to managing climate change impacts.

### Modelling coral eco-evolutionary dynamics

Our model, ReefMod-GBR^21^, simulates individual corals recruiting, growing, competing, reproducing and dying on 3,806 individual reefs interconnected by larval dispersal along the ∼2,300 km length of the Great Barrier Reef (Fig. 1a). Each modelled reef is subject to a specific environmental setting, including water quality, larval connectivity, outbreaks of the coral-eating starfish (*Acanthaster* spp.) and the incidence of cyclones and marine heatwaves. Corals are classified into six groups with variable demographics and susceptibility to disturbances. Temperature-induced mortality (Fig. 1b) is estimated from the Degree Heating Week (DHW; °C-week), a measure of accumulated heat stress exposure^8,22^ proven to be a reliable proxy for coral bleaching and mortality on the Great Barrier Reef^9^. Sensitivity to heat stress is a quantitative trait modelled at the level of individual coral colonies, capturing variations in heat tolerance within^23^ (Fig. 1b, c) and among coral groups^9,21^ (Extended Data Fig. 2c). Heat tolerance is partially inherited^24^, with a heritability coefficient that aligns with empirical estimates for coral thermal traits^25,26^. Incorporating individual-level variations and heritability of heat tolerance enables thermal adaptation across multiple generations. This adaptive process is driven by natural selection in response to successive mass bleaching and mortality events, promoting the survival and propagation of thermally tolerant individuals.

**Fig. 1.**
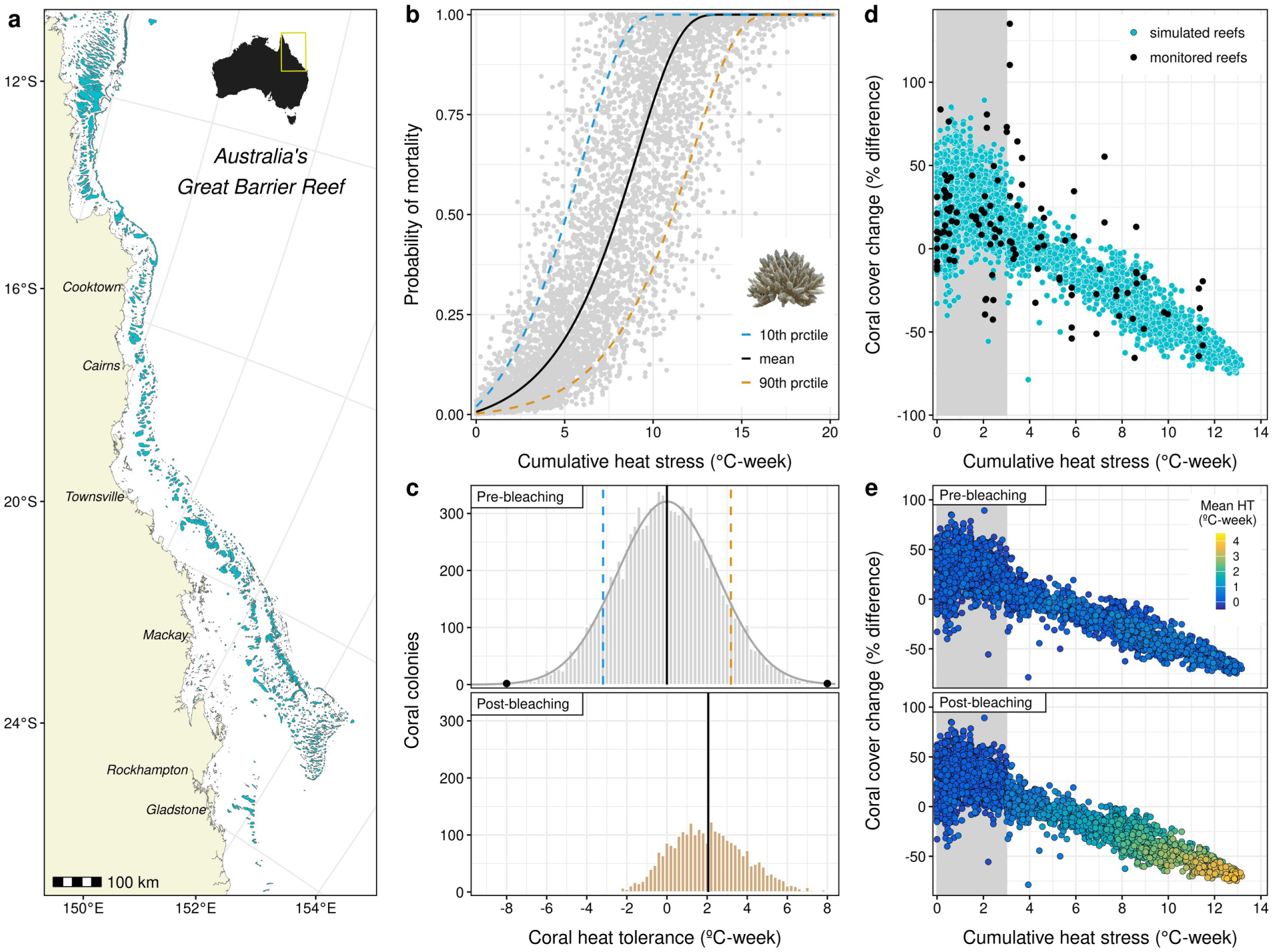
Modelling of coral eco-evolutionary dynamics. **a** Spatial domain of ReefMod-GBR, consisting in 3,806 individual reef units distributed along Australia’s Great Barrier Reef (GBR). **b** Bleaching-induced coral mortality resulting from accumulated heat stress (Degree Heating Week, DHW, in °C-week) at initialisation (i.e., naive response to heat stress). Corals of a given group (here, corymbose acroporids) follow an average dose-response (black curve) modelled from depth-adjusted mortality observed during the 2016 mass bleaching^9,21^. Variability in heat tolerance (±HT; °C-week) among coral individuals (grey dots) is simulated by shifting the dose-response curve horizontally along the DHW axis, reflecting individual deviations from the group mean response at initialisation. The blue and orange mortality curves indicate, respectively, the bleaching response of the 10^th^ and 90^th^ percentiles of the distribution of HT within the group^23^. **c** Simulated HT distributions before and after a hypothetical heat stress of 10°C-week, illustrating the natural selection of heat-tolerant corals. At initialisation, the distribution is truncated normal, with mean 0°C-week (black line), the 10^th^ and 90^th^ percentiles as in **b**, and limits set to ±8°C-week (dots). **d** Impacts of the 2016 mass bleaching on total coral cover (relative change) recorded at 5-10 m depth by independent monitoring surveys^29^ (black dots, N=117 reefs) and predicted by the model (blue dots, N=3,806 reefs), versus reef-level DHW exposure recorded by remote sensing^22^. In the model, bleaching does not occur when reefs are exposed to 0-3°C-week^9^ (grey area). **e** Simulated 2016 bleaching impacts as in **d**, showing changes in the mean tolerance of heat-sensitive coral groups (acroporids and pocilloporids) following selection by bleaching.

We begin by simulating coral community dynamics of the past 15 years to capture the recent selective pressures experienced across the Great Barrier Reef – a cluster of four significant marine heatwaves that occurred in 2016, 2017, 2020 and 2022 (Extended Data Fig. 4a). Starting in 2008 from an average 27% coral cover reported by historical monitoring, simulations reveal large temporal fluctuations across the 3,806 reefs (Extended Data Fig. 5), underlining periods of substantial coral declines as well as a remarkable capacity of recovery. The model predicts an average coral cover of 39% in 2023, reflecting an overall increase from 2008. However, the whole ecosystem experienced prolonged periods of low coral cover as indicated by a 25% annual average. The comparison with monitoring data indicates a credible reconstruction of the past mean coral cover trajectory, with model errors – defined as the annual differences between predictions and observations – being evenly distributed and having a mean difference of –0.3% (SD=± 4.7%). Coral cover reconstructions at individual-reef scales performed reasonably well (Extended Data Fig. 6), despite the challenge of hindcasting patchy disturbances at fine resolution, particularly the damaging waves generated by cyclones^21,27,28^.

The modelling of coral bleaching, parameterised with in situ mortality data from the 2016 marine heatwave^9,21^, allows for an accurate reconstruction of relative coral cover changes observed from independent monitoring^29^ (Fig. 1d). Simulated demographics before and after heat stress predict, on average, a 38% coral decline (interquartile range: 31-45%) to DHW exposures between 8-10°C-week, resulting in an average shift of +1.6°C-week (1.4-1.9°C-week) in the mean heat tolerance of thermally-sensitive taxa (Fig. 1e). Reliable predictions of coral bleaching responses were also obtained in 2017 and 2022, but not in 2020, as no significant mortality was reported despite reef exposure to harmful DHW levels (Extended Data Fig. 7). Since the modelled coral communities were assumed to be relatively naive to heat stress prior to the 2016 mass bleaching, coral mortality decreased during the subsequent bleaching events in 2017, 2020 and 2022, with a mean coral decline of, respectively, 28% (19-39%), 28% (20-37%) and 10% (5-16%) on reefs exposed to 8-10°C-week. Hence, the response to bleaching weakens across successive marine heatwaves due to the progressive elimination of the most thermally-sensitive corals (Extended Data Fig. 8). The strength and persistence of this shift in heat tolerance depend on the intensity of the selective pressure, the rate at which new corals recruit, their thermal traits, and the effective transmission of these traits to subsequent generations.

### Projected heat stress under different warming scenarios

We used daily projections of sea surface temperatures (SST) derived from the Coupled Model Intercomparison Project Phase 6 (CMIP6)^30^ to simulate the impacts of future bleaching events under five alternative GHG emissions scenarios (Shared Socioeconomic Pathway, SSP) assessed in the latest IPCC report^14^. Daily SST projections of ten climate models were downscaled to 10 km using semi-dynamical shelf-sea modelling^31^, allowing us to calculate annual maximum DHW for each individual reef from 2024 to 2100.

Our DHW projections reveal an escalation in the frequency and intensity of marine heatwaves in the coming decades under all emission scenarios (Fig. 2a). Only rapid and drastic reductions of GHG emissions, limiting global warming to +1.5°C (SSP1-1.9), have the potential to reduce SST in the latter half of the century. Adhering to the +2°C limit set by the 2015 Paris Agreement (SSP1-2.6) would only stabilise SST by the end of the century, yet restricting global warming below 2°C requires immediate and strong mitigation actions^32,33^. Under a more likely scenario of ∼2.7°C global warming^6,34^ (SSP2-4.5), SST will continue to rise after mid-century, exposing 50% of the Great Barrier Reef to DHW above 8°C-week (a threshold of widespread bleaching and mortality in heat-sensitive corals^8,35^) at a rate greater than 5 events per decade (Extended Data Fig. 9). Note there is important spatial heterogeneity in the frequency of heat stress, with 10% of reefs expected to experience DHW > 8°C-week only 1-4 times per decade, while 10% would experience it almost annually. Heat stress above 16°C-week, conducive of severe multi-species coral mortality^36^, would affect 50% of the reefs at a rate greater than 3 events per decade. Under more severe warming scenarios (SSP3-7.0: ∼3.6°C; SSP5-8.5: ∼4.4°C; ref. ^5^), heat stress above 16°C-week is projected to affect more than 50% of the Great Barrier Reef at a frequency greater than 6 events per decade after 2060, and annually by 2100.

**Fig. 2.**
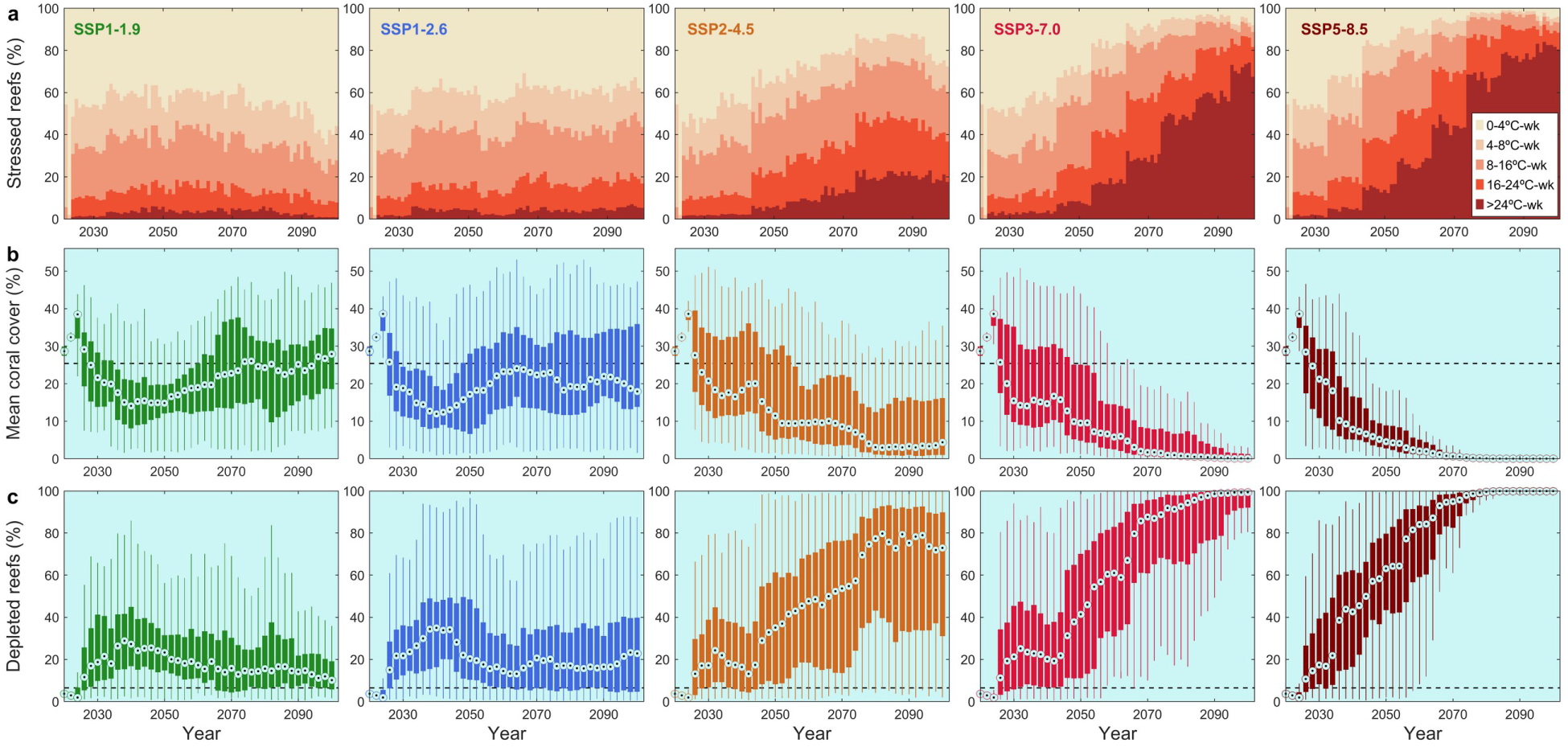
Ensemble projections of Australia’s Great Barrier Reef during 21^st^ century under five climate futures. From left to right, projections are shown for five scenarios of socioeconomic development (Shared Socioeconomic Pathways, SSPs), SSP1-1.9, SSP1-2.6, SSP2-4.5, SSP3-7.0, and SSP5-8.5, leading to a global warming by 2010 of (best estimate^5^) +1.4°C, +1.8°C, +2.7°C, +3.6°C and +4.4°C, respectively. **a** Percentage of reefs (n=3,806) under different categories of heat stress (Degree Heat Weeks, DHW) across all climate projections (n=140 projections for SSP1-1.9, n=200 for all other SSPs). DHW events within each downscaled climate projection were shuffled per decade to generate stochastic fluctuations in the occurrence of bleaching. **b** Biennal likely distributions of the predicted Great Barrier Reef mean total coral cover (percent cover of all corals averaged across all reefs) among 1,000 bootstrap samples, each consisting of 20 individual runs of the eco-evolutionary model drawn at random using the likelihood of the underlying climate projection as weight. Boxes represent the interquartile range (IQR), showing the spread of mean total coral cover predictions between the 25th (Q1) and 75th percentiles (Q3) of the likely distributions. Whiskers extend from each box to the most extreme samples not considered outliers. Outliers (not shown for clarity) are samples greater than Q3 + 1.5 × IQR or less than Q1 – 1.5 × IQR. The central point mark is the median. **c** Distribution of the proportion of reefs with total coral cover < 5%. Boxes, whiskers and central point as for **b**. In **b** and **c**, the dashed lines indicate the hindcast (2008-2023) averages (25.4% total coral cover, 6.5% of reefs in a depleted state).

By comparison, the recent marine heatwaves have exposed 20-30% of the Great Barrier Reef to more than 8°C-week with a maximum of 13.3 °C-week (Extended Data Fig. 4a), although only few reefs were affected by all four events^37^. However, we note that our downscaled ensemble climate model, with SST projections starting effectively in 2014, hindcasts more extreme DHWs than those actually observed (Extended Data Fig. 4b). This discrepancy may reveal a warm bias in our SST projections, potentially overestimating future heat stress events in the early years of climate forecasting.

### Future coral cover trajectories

Predicting the precise timing, location and intensity of acute disturbances (e.g., marine heatwaves, tropical cyclones) is impossible. However, using an ensemble of stochastic simulations, we can project an array of feasible futures to understand differences in reef dynamics under alternative emissions scenarios. Since some CMIP6 climate models are more sensitive than others to anthropogenic climate forcing^38,39^, we designed an ensemble approach that weights climate models based on the likelihood of their equilibrium climate sensitivity (the increase in global temperature resulting from a doubling of atmospheric CO_2_, Extended Data Fig. 10). We assess coral projections based on the ‘likely’ distributions of the average coral cover across the Great Barrier Reef derived annually from the CMIP6 weighted ensemble of climate forecasts.

Our eco-evolutionary simulations project a sharp coral decline over the next 15 years for all emission scenarios (Fig. 2b), with mean coral cover dropping to 14% in 2040 (median of the likely distribution, interquartile range: 8–24%). This corresponds to a 60% decline of the present-day mean coral cover (39% in 2023, Extended Data Fig. 5). Keeping global warming below 2°C (SSP1-1.9: ∼1.4°C; SSP1-2.6: ∼1.8°C; ref. ^5^) would promote coral recovery in the second half of the century as SST ceases to rise, though retrieving the contemporary levels of coral cover would require the most stringent mitigation of GHG emissions (SSP1-1.9). Under a more likely global warming of ∼2.7°C (SSP2-4.5), coral populations would continue to decline, with projections of mean coral cover dropping to 11% (5–29%) by mid-century and 4% (1–16%) at the end of the century. By then, the vast majority of individual reefs (>75%) would be depleted to less than 5% total coral cover (Fig. 2c). Unmitigated GHG emissions (SSP3-7.0 and SSP5-8.5) would drive the Great Barrier Reef to a precipitous decline over the century, achieving a near total loss of coral cover by 2080-2100.

### Scope for natural adaptation

Despite significant differences among SSPs in the severity and frequency of future heatwaves, coral heat tolerance increases at a similar rate across warming scenarios (Extended Data Fig. 11). The maximum rates of thermal adaptation (1.1-1.4°C-week per decade) occur in the first half of the century before gradually decelerating, likely constrained by the approaching limit of heat tolerance (8°C-week). A key indicator of successful adaptation is the persistence of abundant adult populations, particularly among heat-sensitive species that adopt a ‘grow fast, die young’ strategy (Fig. 3a). Under a global warming scenario around 2°C (SSP1-2.6), heat-sensitive corals can adapt effectively, as evidenced by persistent recovery capabilities driven by their fast growth and high reproductive rates. However, with more severe warming (SSP2-4.5, SSP3-7.0), all coral taxa decline at comparable rates (Fig. 3a-b) because their adaptive capacity is insufficient to keep up with the intensifying selection pressure of rising temperatures. Thus, while it can be expected that reefs will eventually be dominated by the most stress-tolerant taxa^7,9,40^, our simulations underline the complexity of projecting coral community composition into the future, as this will result from differences in demography, connectivity, thermal tolerance and rates of adaptation and that have not yet been teased apart^41,42^. Key to coral persistence is the maintenance of coral brood stocks and reproductive potential that promote fast demographic recovery following bleaching. An encouraging finding is that coral juvenile densities would remain near their historical levels if global warming is kept around 2°C (Fig. 3c), thus increasing the scope for adaptation and enhancing demographic resilience throughout the century.

**Fig. 3.**
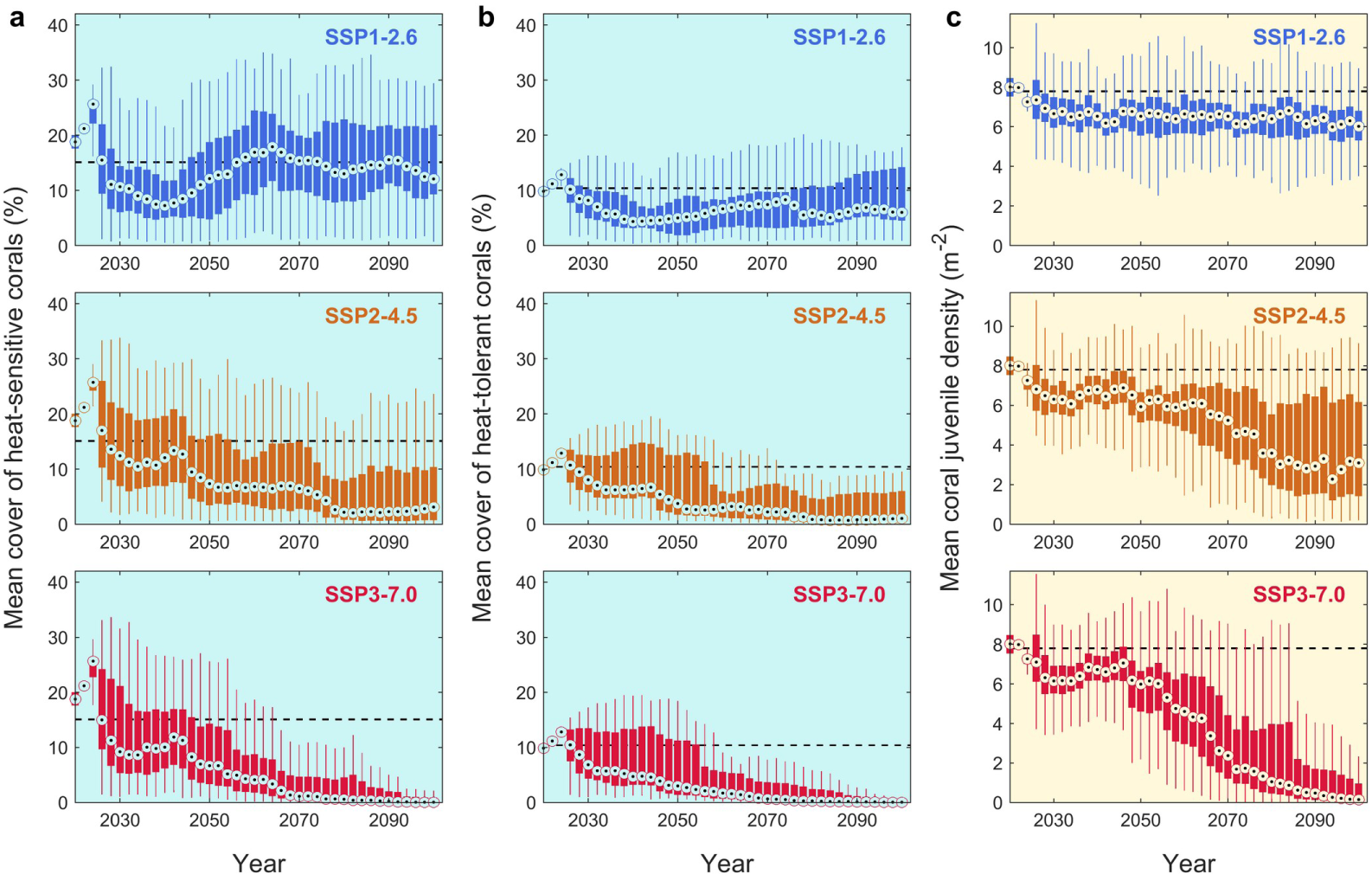
Coral demographics under the three most likely warming futures. **a–b** Ensemble projections of the Great Barrier Reef mean percent cover of heat sensitive (**a**, four taxonomic groups, including acroporids and pocilloporids) and heat tolerant corals (**b**, two taxonomic groups, including large massive and submassive/encrusting corals). **c** Projection of mean density of coral juveniles across all groups. See Fig. 2 legend for the graphical description of distributions.

### Spatial variability of eco-evolutionary responses

Spatial trends in future coral cover exhibit great variability, particularly under relatively modest warming – such as SSP1-2.6 and earlier parts of SSP2-4.5. Mapping the mean state of individual reefs across the weighted ensemble runs (Fig. 4a) reveals large-scale patterns of projected reef health that align with previous studies of thermal refugia and warm spots^43–45^. Thermal refugia were mostly encountered in the southern (Pompey and Swain reef complexes, 20 – 22°S) and far-northern (10 – 12°S) regions, where localised upwelling bring cool deep water to the surface during austral summer^45^. As a result, thermal refugia allow corals to adapt gradually under mild selective pressure while retaining scope for recovery, thereby enhancing long-term coral persistence. However, while different thermal environments result in different rates of evolution, our projections indicate that the availability of thermal refugia will vanish as warming intensifies (SSP2-4.5, SSP3-7.0), leading to fewer reefs maintaining high ecological functioning in the second half of the century.

**Fig. 4.**
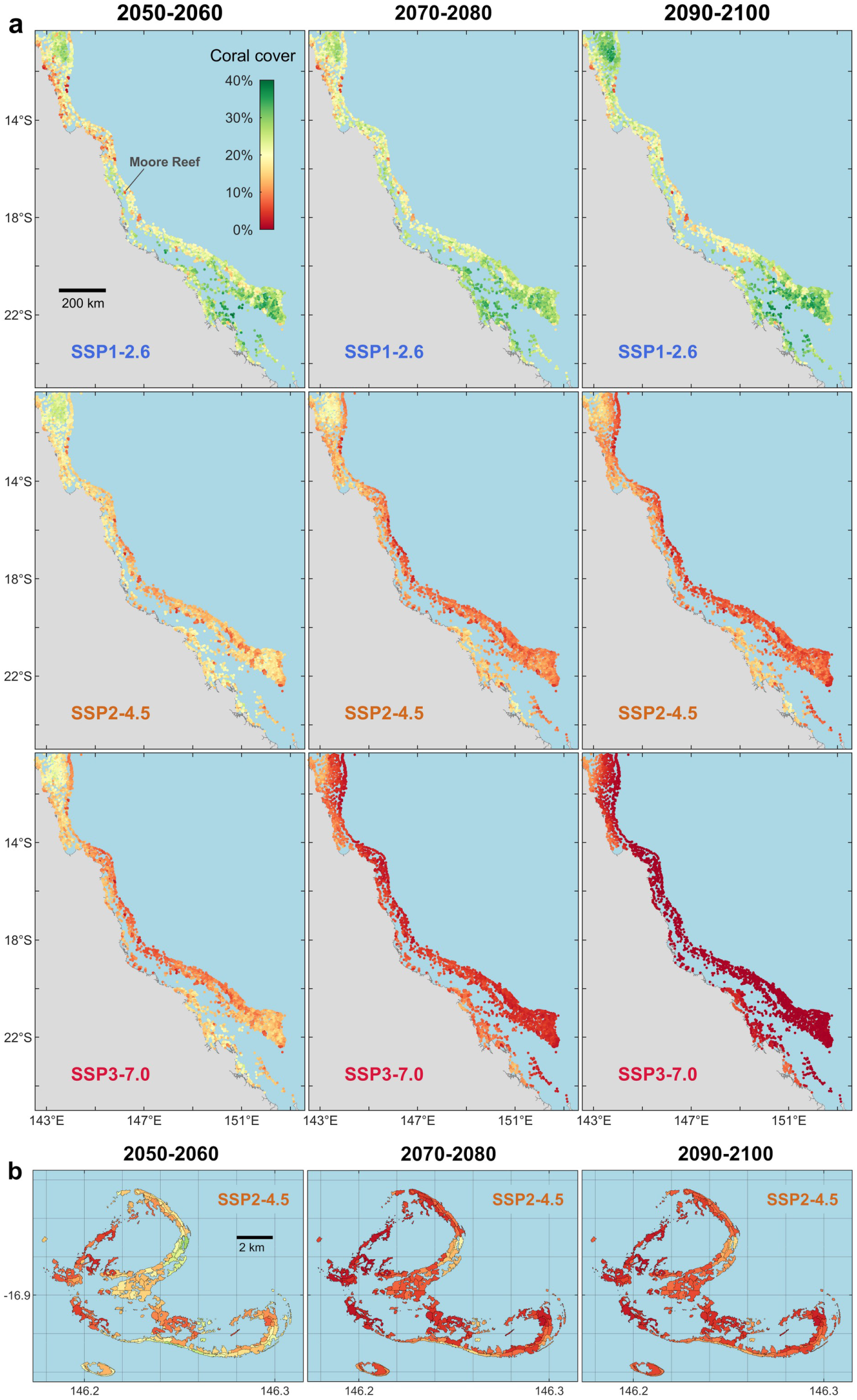
Spatial variability and the progressive disappearance of healthy reefs under severe warming. **a** Annual mean coral cover for 3,806 individual reefs across the Great Barrier Reef over three specific decades under SSP1-2.6 (top), SSP2-4.5 (middle) and SSP3-7.0 (bottom). **b** Annual mean coral cover for 171 reef habitat sites on Moore Reef in the Central Great Barrier Reef (see location in **a**) under SSP2-4.5. Annual means were calculated by averaging the predicted coral cover across all bootstrap samples (1,000 bootstrap samples of 20 individual model runs).

Note that the projected thermal environment was defined at 10-km resolution but thermal refugia may also exist at finer spatial scales (i.e., within individual reefs). Hence, employing a different coral community model^46^ specifically designed to simulate fine-scale (0.5-10 hectares) coral habitats in the Central Great Barrier Reef (Supplementary Text 3), we find that high levels of heterogeneity also exist at finer spatial resolution among the different sites of a reef (Fig 4b). Yet, within-reef variability in coral communities may reduce over time as warming affects corals across all sites.

### Drivers of eco-evolutionary changes

While a simplification of real-world outcomes (Supplementary Text 1), our model incorporates a complex intertwining of environmental drivers, including cyclones, bleaching events, coral-eating starfish outbreaks, and heterogeneous water quality and larval dispersal. To unravel this complexity, we used statistical linear models of likely ecosystem drivers to evaluate which processes exert the strongest influence on future reef state. We analysed this for two timeframes (Fig. 5a): (1) the mid-century, when stress peaks for low emission scenarios, and (2) at the century’s end, when emissions scenarios are most divergent. We find that decreases in the intensity and frequency of coral bleaching – such as those associated with thermal refugia – are most important in promoting total coral cover compared to other disturbances (cyclones, water quality and outbreaks of *Acanthaster* spp.). Indeed, defining thermal refugia and warm spots as representing, respectively, the lower 10^th^ percentile and upper 90^th^ percentile of annual DHW, we find that reefs with higher coral cover are far more common in the cooler refugia (Fig. 5b), especially under mild warming (SSP1-2.6, and SSP2-4.5/SSP3-7.0 by mid-century).

**Fig. 5.**
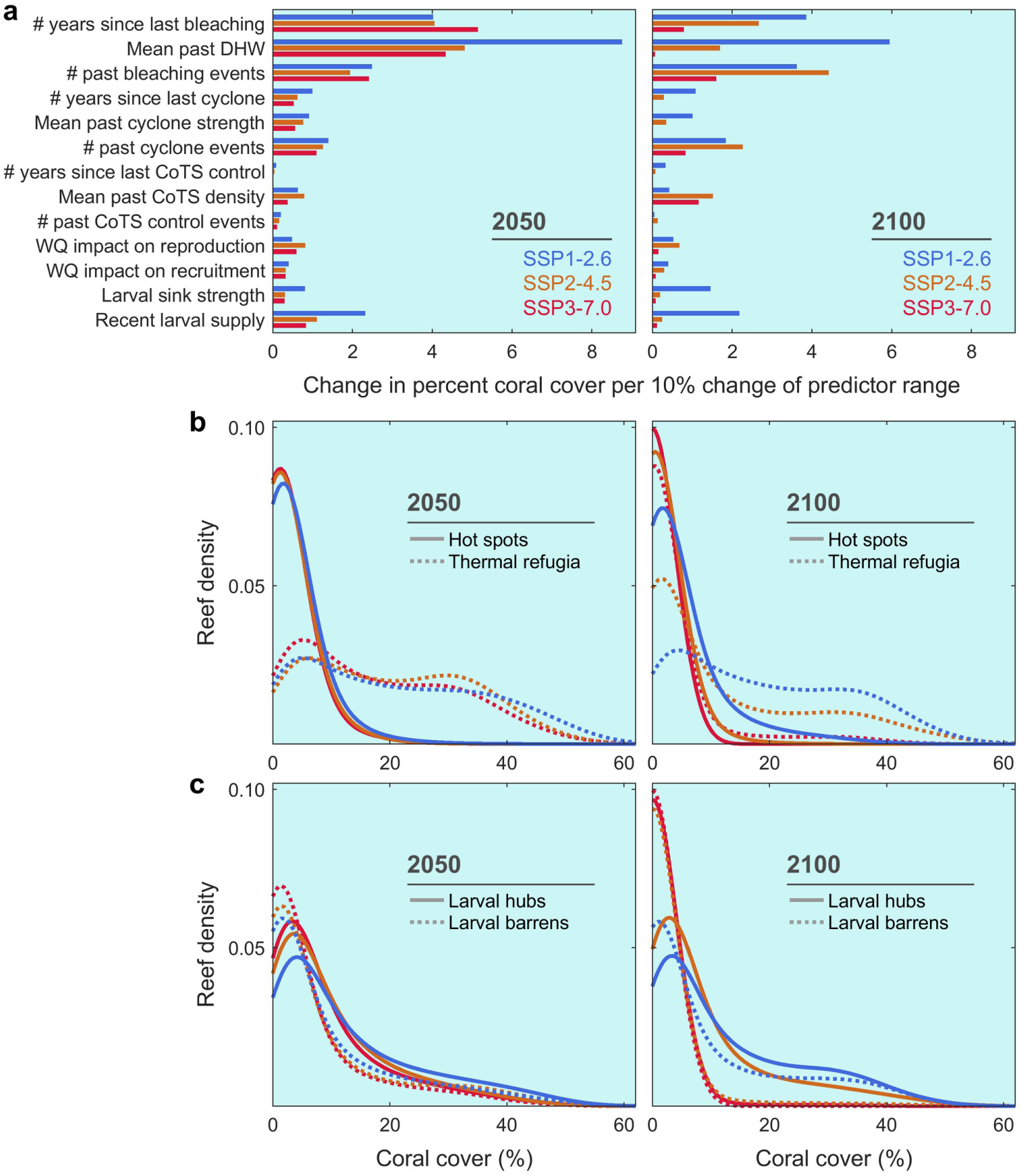
Drivers of future coral persistence under plausible global warming. **a** Factors associated with healthier reef outcomes under SSP1-2.6, SSP2-4.5, and SSP3-7.0 for both mid-century (left) and end-century (right) based on predictions of statistical linear models. The influence of each predictor (Extended Data Table 1) is measured by the change in coral cover (as percentage points) for a 10% change of the predictor within its range. The final row represents external larval supply (i.e., excluding local larval retention). CoTS: crown-of-thorns starfish (*Acanthaster* spp.); WQ: water quality. **b**–**c** Probability (kernel) density distribution of reefs as a function of the reef’s coverage of heat sensitive coral taxa between thermal regimes (**b**, warm spots and thermal refugia) and larval supply regimes (**c**, larval hubs and larval barrens) at both mid-century (left) and end-century (right). Thermal and larval supply regimes are defined, respectively, as those reefs below the 10^th^ or above the 90^th^ percentile for annual DHW and annual amount of external larval supply. Reefs in thermal refugia (i.e., exposed to lowest thermal stress) and in larval hubs (i.e., exposed to highest larval supply) exhibit higher levels of coral cover compared to warm spots and larval barrens. Thermal environments and larval connectivity were calculated as the average heat stress and larval supply experienced in the preceding five years. Larval supply excludes local retention and only captures larvae from external source reefs. Colours of SSPs as in **a**.

We also find a noticeable influence of coral connectivity, either expressed as larval sink strength derived from raw connectivity data or as simulated number of larvae supplied over the past five years (Fig. 5a). This influence is more pronounced under moderate warming (SSP1-2.6), underscoring the compounded effect of heat stress, which diminishes the benefits of larval supply by shortening the periods of recovery. By defining larval barrens/hubs as the lower 10^th^/upper 90^th^ percentiles of the annual amount of external larvae supplied, we observe that higher coral cover is more commonly achieved in larval hubs (Fig. 5c), which is consistent with previous findings using large-scale connectivity^41,42,47^. Moreover, a focus on warm spots shows that reefs with greater access to external larval supply tend to fare better than those with reduced upstream connections, even in the face of high bleaching mortality (Extended Data Fig. 12). Yet, the ability of larval dispersal to predictably benefit coral populations diminishes under higher GHG emissions, particularly towards the end of the century. The disruption to reef state becomes so patchy and severe that larval supply is driven by local retention and sporadic larval dispersions from upstream reefs that retain enough adult corals.

Corals in warm spots and thermal refugia exhibit adaptation to their respective thermal environments. By mid-century, the mean thermal tolerance of corals on refuge reefs is 1-2°C-week lower than that of corals in warm spots (Fig. 6a). As warming worsens, either over time or across SSPs, the range of mean heat tolerance broadens in cooler regions, driven by a higher prevalence of cooler-adapted phenotypes. However, this does not imply that heat-tolerant phenotypes are absent in thermal refugia. In fact, refuge reefs, which support vastly greater coral abundance compared to warm spots (Fig. 5b), have a greater reproductive capacity, allowing them to produce larger quantities of coral offspring, including heat-tolerant phenotypes, though in smaller proportions. The larger volume of larvae means the absolute number of heat-tolerant phenotypes can still be relatively high. Thus, cooler regions are still able to disperse warm-adapted phenotypes to some of the reefs where they are needed.

**Fig. 6.**
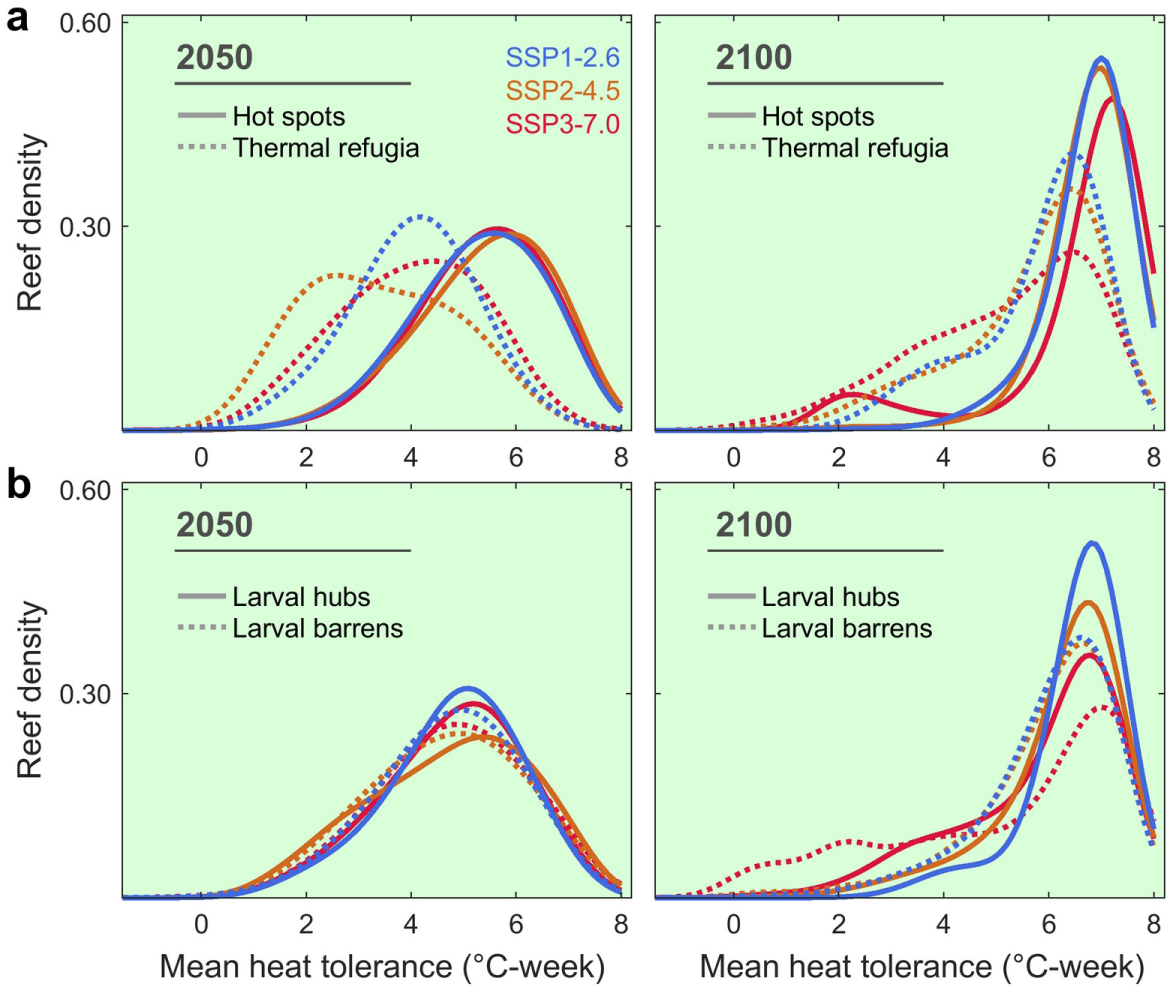
Influence of thermal refugia and larval supply and coral adaptation. **a**–**b** Reef density distribution of mean heat tolerance of heat sensitive coral taxa in the 10^th^ and 90^th^ percentiles of the annual DHW (**a**) and annual amount of external larvae supplied (**b**) for both mid-century (left) and end-century (right). Heat tolerance is defined as the relative accumulated heat stress (± °C-week) a coral colony can withstand compared to the present-day mean response of its taxonomic group (0 °C-week). Recurrent heatwaves progressively shift the mean tolerance on a reef by selecting corals with the highest heat tolerance (within biological limits set to ±8°C-week) over multiple generations. Reefs in thermal refugia slow down the evolution of heat tolerance yet exhibit a greater diversity of heat tolerance values; warm spots achieve greater heat tolerance values at the price of reduced population sizes and lower phenotypic diversity.

Finally, we note that larval barrens and larval hubs display comparable distributions of mean heat tolerance (Fig. 6b). Although connectivity might be expected to impede thermal adaptation with the arrival of maladapted, cooler phenotypes dispersing from thermal refugia^48^, our simulations suggest that larval supply has no apparent influence on local adaptation.

## Discussion

As global warming intensifies, we observe a progressive homogenisation of reef state across multiple spatial scales on the Great Barrier Reef. First, the system-wide average reef state deteriorates, marked by a decline in mean total coral cover in the near term, across all emissions scenarios (Fig 2b). Second, the spatial variability of reef health gradually reduces as the geographical footprint of recurrent marine heatwaves broadens (Fig. 4a). This can be illustrated by observing a single realisation of the future, where the diversity of coral cover trajectories among individual reefs becomes constrained towards the end of century (Extended Data Fig. 1c). Finally, use of a complementary fine-scale ecosystem model parameterised for sites within a small reef cluster finds a progressive loss of coral heterogeneity among sites (Fig. 4b). A loss of heterogeneity across multiple spatial scales could be critically important in ways that are not yet fully understood or incorporated into models. For example, patches of relatively healthy habitat with high coral densities are less likely to experience Allee effects in fertilisation^49^. Their progressive loss will undermine the reproductive success of corals within reefs. Yet, it is important to emphasise that coral communities showed ability to recover from the initial decline under stringent mitigation targets (<2°C, SSP1-1.9 and SSP1-2.6). Perhaps more importantly, coral cover entered a trajectory of recovery when temperatures were still rising (i.e., before mid-century). This is an encouraging finding since it suggests that coral adaptation is possible, provided that the rate of temperature change does not exceed the pace of thermal evolution in corals, which is constrained here by life-history, trait heritability and inter-reef connectivity. This result reaffirms the critical importance of climate policy commitments and the urgency of their implementation to curb the rate of warming before 2050.

Our model incorporates key components of natural selection: trait variation, survival of individuals with advantageous traits, and transgenerational inheritance of trait values. While our parameterisation allows bleaching sensitivity to vary among taxa^40^, we note that the variation amongst thermal phenotypes, the foundation upon which natural selection operates, was parameterised from one of the more sensitive coral taxa (*Acropora* spp.). Specifically, we set an upper limit on heat tolerance which has a notable influence on eco-evolutionary predictions (Extended Data Fig. 13). It is unclear how the scope for thermal adaptation varies across taxa and much remains to be understood about the mechanisms of adaptation itself^50,51^. Moreover, our model does not account for other life history traits that may be affected by rising temperatures^52^ or by the strengthening of thermal resistance^53^, nor does it capture the evolutionary response of symbiont populations to warming, which influence coral growth and heat tolerance^54^. Beyond these simplifying assumptions, we identified a possible warm bias in our downscaled projections of heat stress. Consequently, the predicted global decline in coral cover during the first 15 years of the forecast may be overestimated, although this must be weighed against other model uncertainties, many of our assumptions favouring optimistic outcomes in the simulated coral demographics (Supplementary Text 1). Overall, there was considerable variation in warming projections across the CMIP6 models, leading to a high level of uncertainty around mean coral cover trajectories, both within and between emission scenarios. Given the many uncertainties associated with the response of corals to climate change, we point out that the uncertainty in our model projections increases substantially over time, largely in ways that are difficult to quantify.

The extent to which coral reefs will maintain their function in the coming decades will not only depend on the ability of corals to adapt to rapid warming, but also on our commitment to mitigate GHG emissions and our capacity to implement effective and efficient management strategies. While reducing GHG emissions remains a top priority for the protection of coral reefs, opportunities exist for meaningful management interventions that promote coral adaptation and resilience, even under moderate GHG emissions (SSP2-4.5). By underpinning key mechanisms that drive coral adaptation across a rapidly changing environment, our model can help prioritising multiple interventions, such as protection against local stressors (fishing, coral-eating starfish outbreaks, poor water quality) but also reef rehabilitation^55,56^. To this end, identifying reefs that promote demographic and evolutionary rescue at regional scales is crucial. Protecting thermal refugia may help supporting ecosystem functioning locally while maintaining a greater diversity of phenotypes. Conservation efforts, however, must also include areas under strong selective pressure (i.e., warm spots) to foster genetic adaptation^57–59^. The design of conservation strategies that promote adaptation is in its infancy but one complication is the potential for ‘gene swamping’ where the dispersal of cool-adapted corals into warm spots could slow adaptation^48,57,60^. However, while the potential exists for such disruption on the Great Barrier Reef – where 13% of reefs that constitute refugia from bleaching can disperse corals to 58% of the whole ecosystem^43^ – we found no evidence of a lack of adaptation in thermal refugia and a negligible influence of larval supply on local adaptation. Thus, management strategies for adaptation that encompass multiple thermal environments can be flexible and unconstrained by patterns of larval supply. In fact, larval hubs should be prioritised for building reef resilience since areas with high larval supply are likely to be more resilient even under moderate GHG emissions. Conversely, areas lacking larval supply might be considered for rehabilitation measures. Ultimately, continued investment in reef management is essential even if we move towards lower GHG emissions.

## Methods

### Coral-reef ecosystem modelling

Coral projections under future warming were performed using ReefMod-GBR^21^, a coral individual-based model that simulates coral metacommunity dynamics throughout the ∼2,300 km length of Australia’s Great Barrier Reef (GBR). The model tracks the size of individual coral colonies of six taxonomic/morphological groups under the influence of demographic processes, ecological interactions and disturbances: acroporids (arborescent, plating and corymbose), pocilloporids, large massive corals and submassive/encrusting corals. Coral demography is simulated with a seasonal time step of six months (austral summer: from November to April; austral winter: from May to October) using rates of recruitment, growth, survival and fecundity characteristic of ∼5–10 m deep reef environments. Metacommunity dynamics are captured by tracking the dispersion and settlement of coral larvae throughout a network of 3,806 individual reefs (Extended Data Fig. 1a). Coral stressors include acute disturbances causing coral mortality across the GBR (cyclones, temperature-induced coral bleaching and outbreaks of the crown-of-thorns starfish - CoTS), but also poor water quality (suspended sediment) and unstable coral rubble generated by acute disturbances, which both hinder recruitment success and impede coral recovery. Most processes are stochastic at the colony or reef level, reflecting local variability in demographics and disturbance exposure.

The model has provided credible reconstructions of recent coral cover trends observed between 2008 and 2020 across the GBR^21^. Subsequent applications have explored possible coral futures under climate projections of the fifth phase of the Coupled Model Intercomparison Project (CMIP5) and for different scenarios of reef management^61–63^. Here, we further developed the model with (1) the evolutionary dynamics of coral heat tolerance under selection by heat stress (i.e., mass coral bleaching) and (2) the most up-to-date climate projections (CMIP6) derived from a wider array of greenhouse gas (GHG) emissions. This spatially-explicit, eco-evolutionary model enables us to simulate the potential for coral adaptation under a range of future scenarios of warming across GBR coral communities.

### Model domain and reef definition

An individual reef is represented by a grid lattice of 20×20 cells, each cell approximating 1 m^2^ of the reef substratum. The associated disturbance regime is reconstructed from past exposure (2008-2023) to cyclones, bleaching and suspended sediment concentrations (Extended Data Fig. 1b) inferred at resolutions finer than 10 km (see ref. ^21^ for a detailed description). Here, the area suitable to coral colonisation was refined for each reef individually using high-resolution (10 m) reef mapping derived from remote sensing (Supplementary Text 2). First, an estimate of the 3D surface area of hard reef substrate (down to 20 m depth) was calculated based on reef geomorphological and bathymetry maps^64^; this produced a total area of coral habitat of 13,842 km^2^ for the entire GBR. During simulations, the density of coral and CoTS larvae produced after spawning on the grid lattice of a reef is multiplied by the 3D surface area of that reef to scale up to the amount of larvae produced before dispersal. Second, the proportional area of coral habitat that can be colonised was evaluated based on maps of benthic cover types^64^; this sets a limit to coral colonisation on each individual reef. Coral and CoTS population connectivity among the 3,806 individual reefs was derived from biophysical simulations of coral and CoTS larval dispersal following annual spawning events across the reef network^21,65^.

### Bleaching-induced mortality

Coral mortality caused by bleaching is modelled as a function of accumulated heat stress expressed as Degree Heating Week (DHW, unit: °C-week). Bleaching occurs during the austral summer when DHW ≥ 3°C-week on any given reef^9^ but is prevented if a cyclone also occurs in the season^66^. First, a probability of initial mortality *m* is estimated from a regression model^21^ fitted to GBR observations collected at the peak of the 2016 mass-bleaching event^9^, further extended to account for the sensitivity of multiple taxa at different depths:

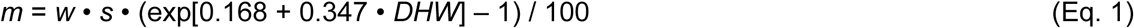

where *w* is a depth-related attenuation coefficient of heat stress and *s* is a sensitivity coefficient for specific taxa. With *w* = 1 and *s* = 1, Eq. 1 predicts the community-level response recorded at 2 m depth. Coefficient *w* is modelled (Extended Data Fig. 2a) using observations of bleaching mortality collected in the Northern GBR along a depth profile of light attenuation^67^:

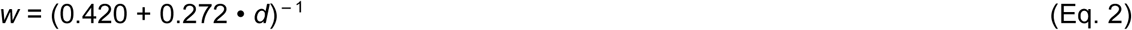

We modelled reefs at an average depth of 7 m, considering the modelled coral demographics are representative of 5–10 m forereef environments, thus effectively fixing *w* to 0.43. The sensitivity coefficient *s* was determined based on observed taxon-specific responses to bleaching^9^ and ranges between 0.25–1.70 (ref. ^21^).

In a second step, initial mortality *m* is capped to 1 and extended to include subsequent mortality occurring after the peak of the bleaching event. A previous calibration^21^ found that the long-term (8 months) losses of percent coral cover reported on the GBR following the 2016 mass bleaching^9^ can be reproduced by simulation with the following transformation:

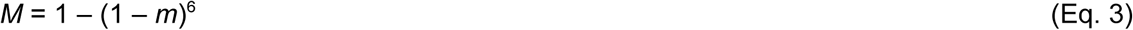

where *M* is the probability of total mortality within each taxonomic group. This transformation confers a sigmoid shape to the DHW/mortality relationship (Extended Data Fig. 2b) which sets 50% coral mortality at ∼8°C-week and ∼13°C-week for heat sensitive (the three acroporid groups and pocilloporids) and heat tolerant taxa (large massive and submassive/encrusting corals), respectively (Extended Data Fig. 2c).

### Heat tolerance of individual corals

Significant variation in heat tolerance can occur among corals of the same species within a specific thermal environment^23,68,69^. To capture this variability within a taxonomic group, we define the heat tolerance (HT) of a coral colony as a quantitative trait that reflects its relative ability to survive heat stress^23^ compared to the group mean survival rate, i.e., the representative DHW/mortality relationship of the group at initialisation. Specifically, HT measures the difference in the amount of accumulated heat stress (± °C-week) that a coral colony can withstand compared to the group mean response:

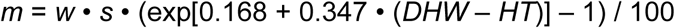

This essentially shifts the curve of initial mortality along the axis of DHW values: a positive and negative HT value confers lower and higher bleaching mortality, respectively, compared to the group mean for the same DHW value. It can be re-written as:

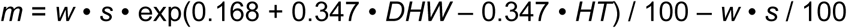

The term *w* • *s* / 100 being negligible (< 0.01, less than 1% mortality), a fair approximation is:

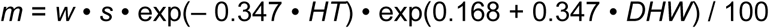

Hence, the specific bleaching response of a coral colony can be obtained from Eq. 1 using a new sensitivity coefficient *s\** expressed as:

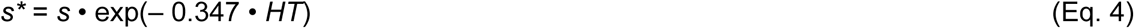

We assume that HT values at initialisation within a taxonomic group follow a normal distribution of mean 0, which corresponds to the average bleaching sensitivity (s) of that group. To get a realistic estimate of the variability of HT, we used the results of a heat stress experiment performed on the corymbose *Acropora digitifera* collected on a shallow outer reef crest in Palau, Western Pacific Ocean^23^. Coral fragments were exposed to an increasing temperature and assessed for signs of heat stress, allowing the modelling of a 0–1 index of bleaching and mortality (BMI) as a function of daily DHW values. As a result, there was a difference of 4.8°C-week (3.1–6.8°C-week 95% confidence interval) between the 10th and 90th percentiles of the population measured at the cut-off BMI level of 0.12 (i.e., 12% mortality). Assuming the BMI is a functional equivalent to *M*, we infer that this difference is ∼6.4°C-week at 100% mortality (Extended Data Fig. 3a). For a standard normal distribution, the 90^th^ percentile has a score of ∼1.28. Therefore, a normal distribution of mean zero and a 90^th^ percentile score of 6.4/2 has a standard deviation of 6.4/(2×1.28)=2.5. We thus assume that present-day HT values follow a normal distribution of mean zero and standard deviation σ=2.5 within each taxonomic group. At initialisation, a value is drawn from this distribution (Extended Data Fig. 3b) for each created coral. Corals retain their assigned HT throughout their lifetime. When bleaching occurs, a probability of mortality is calculated for every colony following Eq. 1–4, and survivors are determined by binomial draws using the associated probability.

Because a normal distribution has tails extending to infinity, negative and positive limits to HT (±HT_max_) are required to avoid creating unrealistic heat tolerance values. A conservative range would set limits to the bottom and top 1% of the empirical distribution, which correspond approximately to −6 and +6°C-week, respectively (Extended Data Fig. 3b). We extend this range by 2°C-week on each side of the distribution (i.e., |HT_max_|= 8°C-week) considering that: (i) this distribution draws from experimental data gathered for a single species; a broader range of HT values can be legitimately assumed when inferring for multiple taxa; (ii) extreme thermal phenotypes might emerge from complex interactions among mechanisms of selection, acclimation and genetic mutations.

### Cross-generational heritability of heat tolerance

Thermal adaptation occurs as a trans-generational response to natural selection driven by mass bleaching and mortality events. Inheritance of heat tolerance is modelled based on the narrow-sense heritability (*h*^2^) which measures the proportion of variation in this trait that is attributable to additive genetic effects (i.e., those genetic effects that are inherited, as opposed to dominance and epistasis)^70,71^. As such, heritability expresses the similarity of thermal phenotypes between parents and offspring, and being a proportion, is confined from 0 (offspring and parental phenotypes are unrelated) to 1 (perfect transmission of parental phenotypes).

Based on empirical estimates^25,26^, we set *h*^2^=0.3 for the heritability of heat tolerance and assume it constant in space and time (but see ref. ^70^). To determine thermal phenotypes (i.e., HT) in coral offspring, we first predict the mean heat tolerance of the new generation from the selected parents following the univariate breeder’s equation^72,73^:

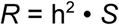

where *R* is the per generation response to selection expressed as the change in mean heat tolerance in the next generation and *S* is the selection differential (i.e., the change in mean heat tolerance after selection within the parental population). Specifically, *S* (°C-week) within each taxonomic group is estimated at every time step t as:

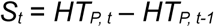

where *HT_P_* is the mean heat tolerance of adult colonies weighted by the amount of their reproductive outputs (function of colony size as per ref. ^21^), such that colonies with greater fecundity contribute more to the mean parental phenotype. Assuming the offspring phenotypes (*HT_O_*) are normally distributed, their mean value is obtained by adding the generational response to selection to the mean parental phenotype *before* selection:

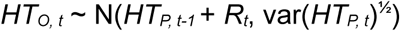

Note the variance of heat tolerance in the population of offspring is assumed to match the fecundity-weighted variance of parental phenotypes *after* selection, based on the expectation that phenotypic variance is reduced under strong selective pressure. In the particular case where no gravid colonies were present at t-1, *HT_P, t-1_* was set to 0 so that the selection differential is calculated relative to the present-day distribution of heat tolerance.

Since corals produce millions of offspring, the computation is only performed for a representative sample of 1,000 offspring HT phenotypes per taxonomic group per reef. The sampled HT phenotypes are then dispatched across the reef network according to probabilities of larval dispersal specified in connectivity matrices. Practically, a pool of 1,000 incoming HT phenotypes is created for any given sink reef by sampling within each pool of outgoing HT phenotypes (i.e., at the reefs of origin) proportionally to the associated source-sink dispersal probability. Hence, the distribution of HT phenotypes within a pool of incoming larvae reflects the contribution of each source reef to larval supply^74^ A coral recruit is then assigned one of the incoming HT phenotypes through random sampling (with replacement).

### Historical regime of environmental forcing

The model first simulates the recent (2008-2023) dynamics of coral and CoTS populations from temporally- and spatially-realistic environmental regimes^21^ (Extended Data Fig. 1b). To ensure model simulations accurately reflect real-world conditions, the cover of each coral taxonomic group at each individual reef is initialised based on historical observations from monitoring data (see details in ref. ^21^). As in previous applications of the model, coral and CoTS larval connectivity (i.e., source-sink dispersal probability) is derived from biophysical simulations of larval dispersal^65,75^ performed over six spawning seasons (water year 2010, 2011, 2012, 2014, 2015 and 2016) using a three-dimensional hydrodynamic model of the GBR^76^. For corals, larval particles were released at actual spawning dates following historical observations of mass coral spawning across the GBR, resulting in a total of 25 simulated spawning events^75^. For CoTS, four spawning events were simulated for each spawning season^65^. For both coral and CoTS, connectivity matrices from multiple spawning events were combined to create a single matrix for each of the six water years (see details in refs ^65,75^). The resulting annual connectivity matrices were applied as a recursive chronological sequence over the 2008-2023 period^21^. Past exposure to cyclones was reconstructed from sea-state predictions of wave height^27^ paired with estimates of cyclone category inferred from the closest distance to actual cyclone tracks^77^. Historical heat stress was determined from satellite-derived records of maximum annual DHW at 5-km resolution from NOAA Coral Reef Watch^22^. Suspended sediment concentrations were inferred from 2010-2018 simulations of the eReefs coupled hydrodynamic-biogeochemical model^76,78^. CoTS management was simulated by culling CoTS populations on a per reef basis^61^ following the CoTS Control Program implemented across the GBR Marine Park^79^. Hindcast coral cover reconstructions between 2008-2023 were simulated by repeating 20 stochastic runs of the historical environmental regime. In each run, normally-distributed noise was injected into the mechanisms of population initialisation, recruitment and mortality events across individual reefs. This process generates demographic fluctuations reflecting the natural variability of reef populations.

### Forecast scenarios of environmental forcing

Scenarios of future heat stress during the 21^st^ century were developed using an ensemble of coupled atmosphere-ocean general circulation models (AOGCMs) of the Coupled Model Intercomparison Project Phase 6 (CMIP6) database^30^. AOGCMs simulate possible future climates at regional scales under the Shared Socioeconomic Pathway (SSP) framework, a set of alternative scenarios of societal and economic development leading to different trajectories of atmospheric carbon concentrations^80^. We extracted climate projections simulated daily between 2014 and 2100 by ten AOGCMs (Supplementary Table 1) under five scenarios of GHG emission entailed in the last Intergovernmental Panel on Climate Change (IPCC) Assessment Report^5^ (SSP1-1.9, SSP1-2.6, SSP2-4.5, SSP3-7.0, SSP5-8.5). SSP1-1.9 and SSP1-2.6 are two scenarios of strong mitigation, in which global warming by 2100 is limited to 1.5°C and 2.0°C above preindustrial (1850-1900) levels in line with the 2015 Paris Agreement targets. SSP2-4.5 is an intermediate scenario in which GHG emissions remain around contemporary levels until the middle of the century, with end-of-century warming estimates around 2.7°C above preindustrial temperatures. SSP3-7.0 and SSP5-8.5 are non-mitigation scenarios with atmospheric carbon concentrations doubling from current levels by 2100 and 2050, respectively, leading to global warming around 3.6°C and 4.4°C.

As a result, an ensemble of 47 climate realisations were available at coarse spatial resolution (80-500 km), with SSP1-1.9 being only available for seven AOGCMs (Supplementary Table 1). Each daily warming projection was downscaled to 10 km using semi-dynamical shelf-sea modelling across the 0-50 m depth range^31,81^. Briefly, surface-level atmospheric variables predicted by AOGCMs were combined with high-resolution bathymetry and tidal forcing to simulate daily sea surface temperatures (SST) based on local physical properties of the water column. For each downscaled SST projection, DHWs were calculated as the daily summation of SST anomalies exceeding 1°C above the maximum monthly mean temperature (MMM) over a 12-week running window, following ref. ^22^. The MMM of each 10-km pixel was calculated over the period 1985-2012 from the NOAA Coral Reef Watch climatology^22^. Annual maximum DHWs were calculated over an “austral year” (August to July) to avoid the double counting of bleaching events across two calendar years during austral summers, and assigned to each of the 3,806 reefs based on the closest 10-km pixel.

Because only one warming realisation was available for each AOGCM/SSP, the resulting future for any given reef is one series of heat stress events occurring at specific times (i.e., years). To allow for fluctuations in the timing of future coral bleaching, we generated 20 stochastic sequences of heat stress between 2024–2100 for each warming scenario. This was achieved by randomly sampling, without replacement (i.e., shuffling), the DHWs predicted for all reefs within each decade, generating an ensemble of 940 stochastic warming futures. By shuffling the DHW years for all 3,806 reefs together within each decade individually, we ensured that the spatial distribution of heat stress was preserved for any given DHW year, while maintaining the long-term warming trend forecasted by each climate model.

Each of the 20 stochastic DHW futures of a given AOGCM/SSP was paired with one of the 20 stochastic hindcast reconstructions and one specific timeseries of 20 stochastic future (2024–2100) of cyclone tracks^82^. Applying the same scenarios of future cyclones with each AOGCM/SSP ensures consistency in the comparison of warming scenarios. Hindcast spatial distributions of suspended sediments and coral/CoTS larval connectivity were applied in recursive sequences until the end of the century (Extended Data Fig. 1b).

### Ecosystem projection analysis

A key uncertainty in projections of future climate change is the sensitivity of global warming to GHG emissions^83,84^. Equilibrium Climate Sensitivity (ECS), which quantifies the increase in global temperature resulting from a sustained doubling of CO_2_ concentrations, is a standardised measure for comparing the sensitivity of different AOGCMs to changes in greenhouse gas forcing^85^. While our selection of AOGCMs covers a broad spectrum of ECS values (2.6 – 5.3°C, Supplementary Table 1), caution has been raised over the selection of models associated to unrealistically high ECS^38,39^. Equilibrium climate sensitivity is very likely (90% probability) to lie between 2.3 – 4.7°C (ref. ^86^), yet a significant number of CMIP6 models have an ECS value above this range^38,85^. While the warming projections of these ‘hot’ models^39^ are biased towards the high end, they cannot be ruled out as implausible^85,86^.

To avoid biased projections favoring excessive warming, we assessed future reef health and associated uncertainty by assigning weights to each eco-evolutionary simulation based on the likelihood of the ECS of the underlying AOGCM. First, we inferred the relative likelihood of each AOGCM (Extended Data Fig. 10) given its ECS value based on a Bayesian-derived probability distribution of Earth’s climate sensitivity^86^. Then, we employed bootstrap sampling within each SSP ensemble of simulations (140 runs for SSP1-1.9, 200 runs for all other SSPs) to randomly select eco-evolutionary runs. In this process, a bootstrap sample for a given SSP comprises 20 runs of the model, one for each scenario of future cyclones paired with one sequence of DHW drawn from the available AOGCMs using the associated likelihood as the weighting factor. This approach ensures that (1) cyclone variability is equally represented across all bootstrap samples, (2) final projections are constrained by the most realistic warming limits and (3) all AOGCMs are considered for a reliable assessment of uncertainty around future warming. A total of 1,000 bootstrap samples were constructed for each SSP scenario.

For each individual run, percentage total coral cover, density of coral juveniles (defined as coral diameter between 1-5 cm) and thermal tolerance for each coral taxonomic group was averaged across the 3,806 individual reefs with the log-transformed area of each reef as weight. For every SSP ensemble, these ecological metrics can be tracked through a distribution of 20,000 estimates (20 runs × 1,000 bootstrap samples) obtained at yearly intervals from 2008 to 2100.

### Drivers of ecosystem changes

In any given simulation some reefs fare better than others. Our goal here was to identify the drivers of success and compare them among GHG emissions pathways and between mid- and late century. For each AOGCM and SSP, we took 20 independent simulations of the dynamics of all 3806 reefs until 2100. This resulted in between 532,840 and 761,200 individual reef trajectories per SSP, depending on the number of AOGCMs available. Two census dates were created, which included: 2050 (for mid-century), when stress levels approach maximum for the low emissions scenarios, and 2100 (for late century), when stress levels diverge greatly among SSPs. We then took all simulations for a given time period (e.g., mid-century) and normalised the predictors and response (total coral cover per reef) by z-score. Responses were normalised to ignore differences in the absolute cover between census years. Predictors were normalised because of diversity in their units of measurement (Extended Data Table 1). The only exception was the cumulative number of larvae arriving at reefs in the last five years due to high variability in z-scores and the need to use the log of larval supply to stabilise variance.

The role of predictors on future reef state was evaluated using simple linear models^87^. We did not include random effects since we did not intend to create a model for future prediction and their incorporation could mask desirable insights. For example, a random (or fixed) effect might help explain why reefs under one climate model fared better, on average, than those under another climate model, but this is a distraction from our perspective; our primary interest is that healthier reefs were associated with, e.g., less frequent bleaching events, which happened to occur less often in some models than others. We do not provide estimates of predictor significance because the data result from models and any variable can become significant with enough simulations.

Predictors of relatively healthy reefs were identified using two methods. First, the explanatory power of the transformed predictor in the linear model, represented by the partial coefficient of determination^88^ (R^2^). The second was more complex because we wished to provide a comparable measure of the impact of each variable on reef state, recognising that most predictors are measured on different scales. To do this we first rescaled the model coefficient of each predictor so that it reflected units of the original predictor’s impact on coral cover. In order to provide some parity among predictors with different units, we rescaled the effect size so that it measured the change in coral cover associated with a 10% increase of the predictor across its range. For example, a result of 4% cover associated with metric X implies that a 10% increase within the range of metric X would increase average coral cover by 4 units of cover, with all else being equal.

### Sensitivity analysis

We assessed the sensitivity of future coral cover predictions to changes in key input parameters, focusing on those likely to have the greatest influence on coral eco-evolutionary simulations (Extended Data Table 2): susceptibility to heat stress (two parameters), thermal adaptation (2), recovery potential (3), larval supply (3) and susceptibility to cyclones and CoTS outbreaks (3). Using a baseline scenario of future cyclones (run #1) across three warming projections (SSP1-2.6, SSP2-4.5, SSP3-7.0) from the MIROC-6 climate model (supplementary Table 1), we systematically varied each parameter by ± 20% from its default value, while keeping all other parameters at their default settings. The effect of these parameter changes was assessed in terms of their impact on the GBR mean coral cover projected over the simulated period (Extended Data Fig. 13).

Among the parameters of coral recovery (colony growth, fecundity, larval supply), growth rate and the maximum density of coral settlers (α.coral) were the most influential, causing variations in the GBR mean coral cover of up to ±5% (percentage points) under moderate warming. However, their influence diminishes as warming intensifies. Parameters related to exposure and sensitivity to heat stress (depth-attenuation of DHW and standard deviation of the naive distribution of heat tolerance) also had a substantial effect, with variations up to ±2% on the projected GBR mean coral cover under moderate warming, though their impact also decreased with more intense warming. Heat tolerance limits had a notable effect under SSP1-2.6 by constraining coral adaptation in the later half of the century. In particular, expanding the tolerance limit to ±9.6°C-week would improve the mean coral cover by almost 5%, though the difference from the baseline scenario shrinks towards the end of the century. The influence of heat tolerance limits is considerably reduced under more intense warming, underscoring the dominant impact of escalating heat stress beyond coral adaptive capacity. Conversely, variations in the heritability of heat tolerance have no noticeable effects in any of the scenarios. Parameters controlling the magnitude impacts of cyclones and CoTS have relatively limited impacts.

## Data availability

Data used and produced by this study are available at: https://github.com/ymbozec/GBR_futures

## Code availability

Code of the model version of ReefMod-GBR that has been developed and used for this analysis (REEFMOD.7.0_GBR) is publicly available at: https://github.com/ymbozec/REEFMOD.7.0_GBR

Codes used for all figures and analyses are publicly available at: https://github.com/ymbozec/GBR_futures

## Acknowledgements

We acknowledge the Traditional Owners of the Great Barrier Reef, and pay our respects to their Elders past, present, and emerging and recognise their continuing spiritual connection to sea country. This work was funded by the Reef Restoration and Adaptation Program, a partnership between the Australian Government’s Reef Trust and the Great Barrier Reef Foundation. We thank D. Mead, C. Robillot, M. Baskett, C. Kenkel, A. Baker and C. Castro-Sanguino for insightful comments at various stages of this work. We thank J.T. Morris from NOAA for a thoughtful review of the manuscript. J. Gilmour, M. Gonzalez-Rivero, D. Barneche, B. Robson and J. Moore contributed to the development of the model C∼scape, and R. Ferrari, C. Doropoulos, M. Toor, S. Gordon, K. Crossman and M. Puotinen contributed to its parameterisation.

## Author contributions

Y.-M.B. and P.J.M. conceived the study, performed the analysis and wrote the first draft of the manuscript. Y.-M.B. designed an implemented the model ReefMod. J.K.M., B.A.N. and P.H. downscaled the warming projections. A.K.C., V.H.-B. and J.-C.O. designed and implemented the model C∼scape. All authors contributed to interpreting the results and improving the paper.

## Material and correspondence

Correspondence and requests for materials should be addressed to Y.-M. Bozec.

## Extended data

**Extended Data Fig. 1.**
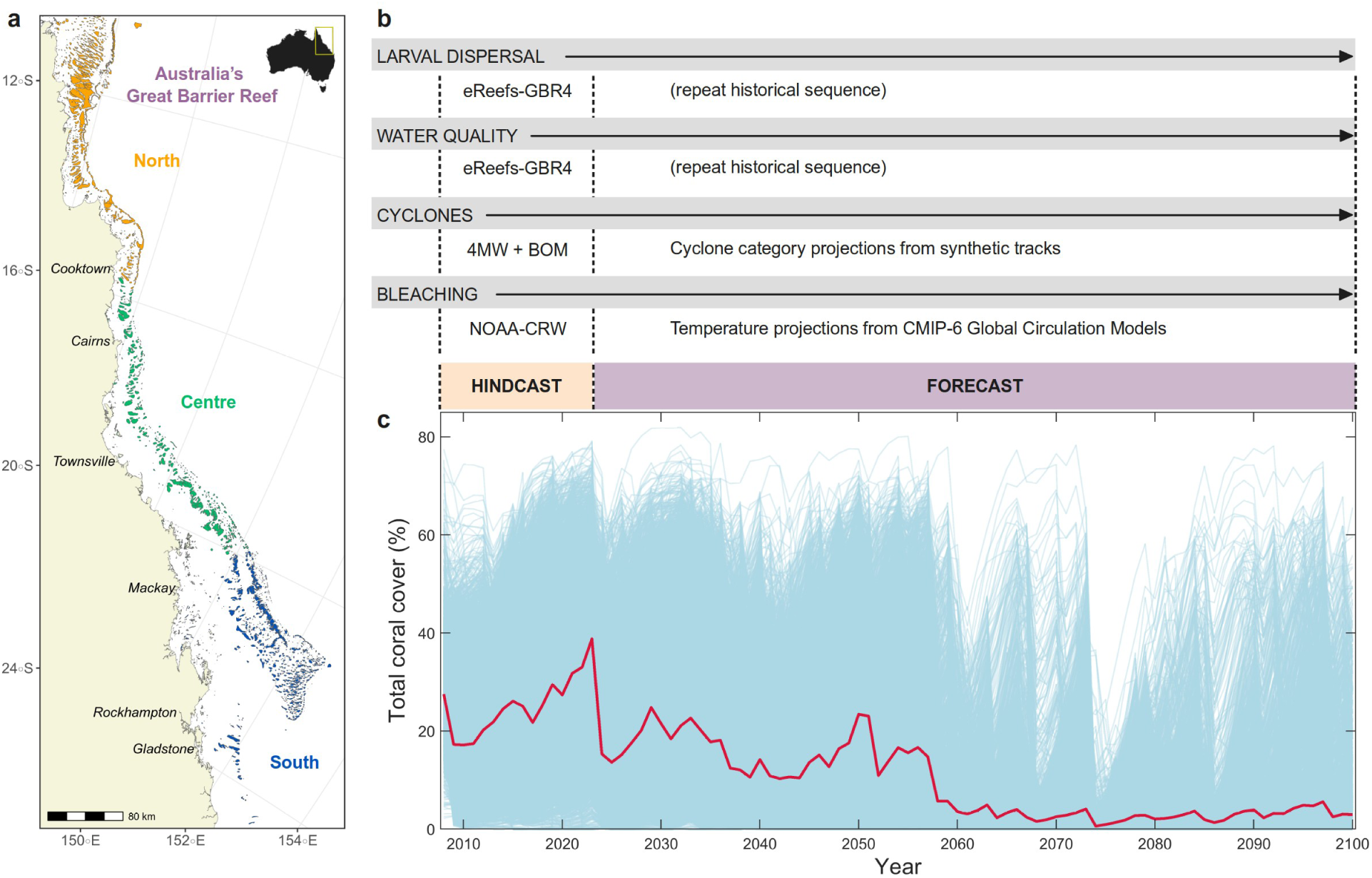
Model domain and reef definition of ReefMod-GBR. **a** Spatial domain of the model consisting in 3,806 individual reefs distributed along Australia’s Great Barrier Reef (GBR). **b**, environmental drivers of coral metacommunity dynamics used for spatially-realistic reconstructions^21^ (hindcast) and future projections (forecast). Coral and CoTS larval connectivity is derived from biophysical simulations following six spawning events^65,75^ using a three-dimensional hydrodynamic model of the GBR^76^ at 4-km resolution (eReefs-GBR4). Water quality is represented by suspended sediment and chlorophyll concentrations inferred from eReefs-GBR4 coupled hydrodynamic-biogeochemical simulations^76,78^. Past exposure to cyclones is reconstructed from predictions of damaging wave height using the model 4MW^27^ combined with estimates of cyclone category from historical tracks^77^. Projection of cyclone categories is determined from a library of future cyclone tracks for the GBR^82^. Historical heat stress is determined from past records of maximum annual DHW at 5-km resolution^22^. Future heat stress is informed by CMIP6 climate model simulations downscaled at 10-km resolution under five emission scenarios. **c**, a single realisation of the future (here, one run using DHW projections from the climate model CNRM-ESM2-1 under SSP2-4.5) shows the variability among individual reef coral cover trajectories (blue lines). The red line indicates the average coral cover weighted by the area of each individual reef.

**Extended Data Fig. 2.**
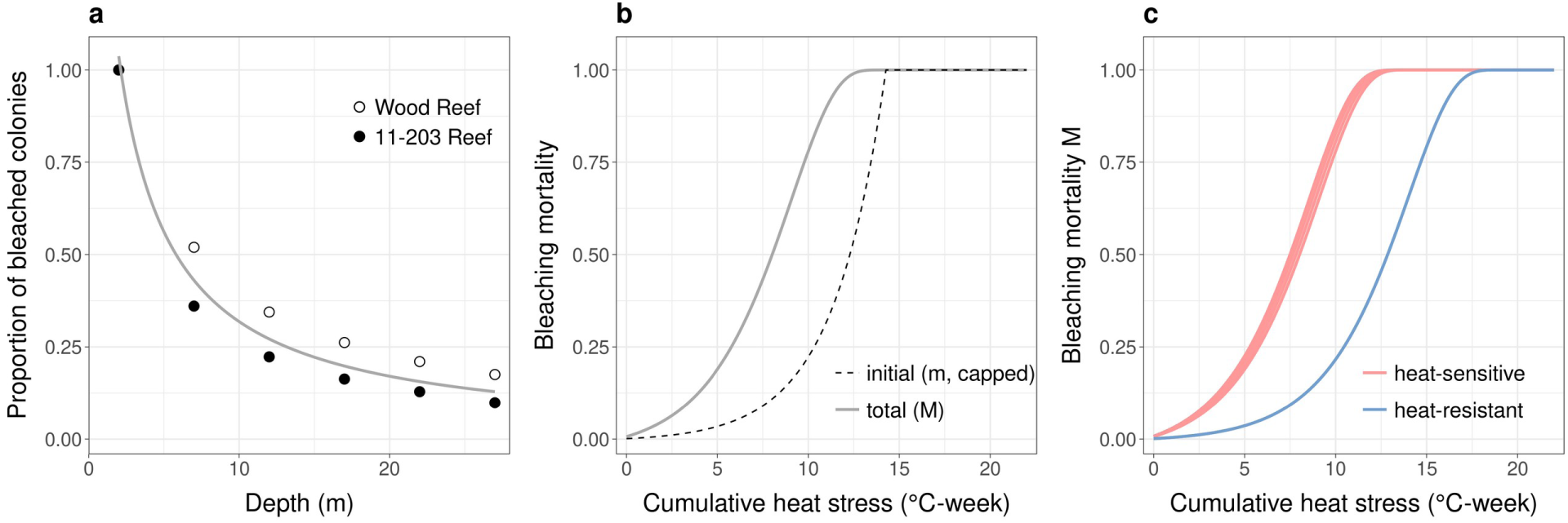
Modelling of bleaching-induced mortality of corals. **a**, modelling of bleaching prevalence along a depth profile of light attenuation as observed in the Northern GBR four weeks after the peak of a mass-bleaching event^67^. Bleaching prevalence was measured at Wood Reef (DHW = 9.0 °C-week) and 11-203 Reef (DHW = 5.8 °C-week) and re-expressed as the proportion of bleached colonies relative to that observed at 2 m depth to be fitted with a linear model after an inverse transformation (R^2^ = 0.78). **b**, empirical relationship between cumulative heat stress (*DHW*) and initial mortality (*m*, Eq. 1) and the resulting long-term, total mortality (*M*, Eq. 3) for corymbose acroporids extrapolated at 7 m depth. **c**, total mortality along the gradient of cumulative heat stress at 7 m depth for the six coral groups, showing differences in mortality among taxa due to variable bleaching susceptibilities (sensitivity coefficient *s*, Eq. 1). For the three acroporid groups and pocilloporids (heat-sensitive taxa, red lines), *s* was set to 1.5, 1.6, 1.4, and 1.7, respectively. For submassive/encrusting corals and large massive corals (heat-tolerant taxa, blue lines), *s* was set to 0.25. Sensitivity coefficients were determined based on taxon-specific mortalities observed on the GBR at the peak of the 2016 mass bleaching^9^.

**Extended Data Fig. 3.**
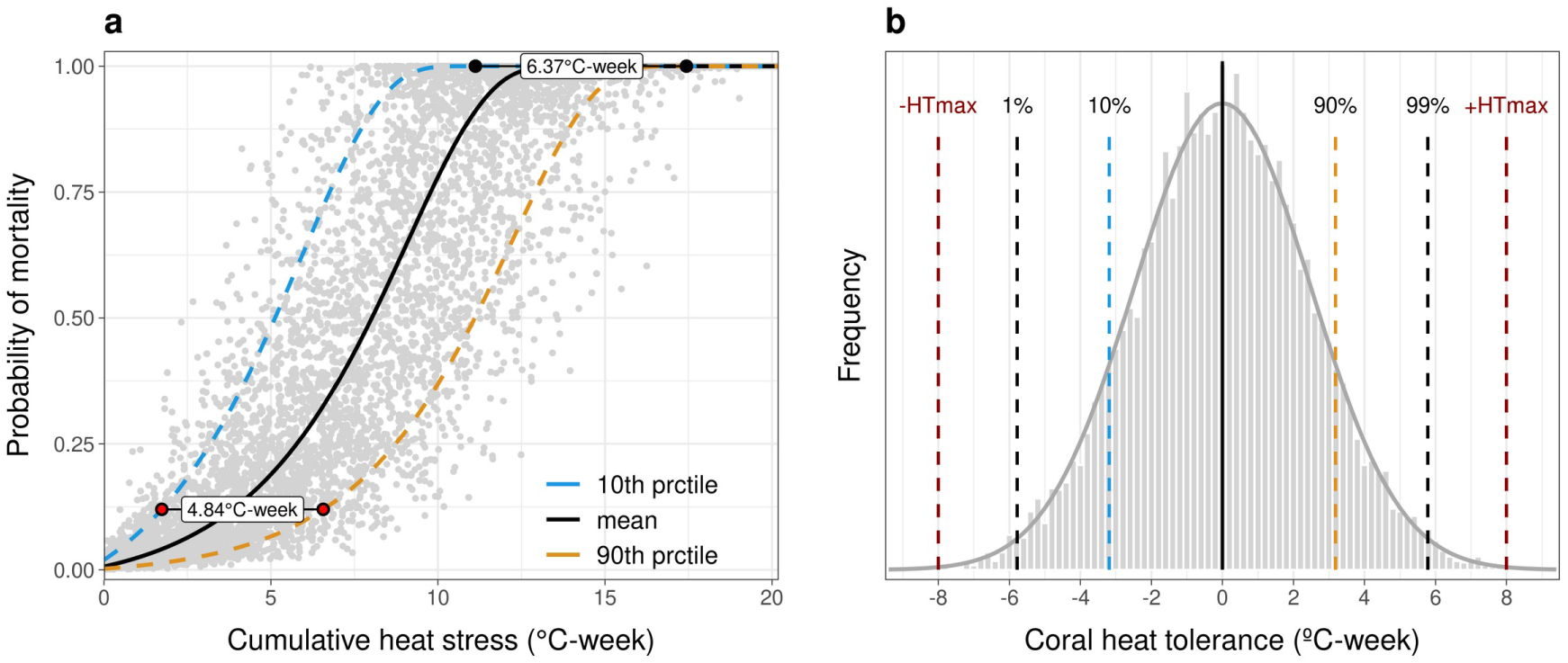
Modelled individual variability of heat tolerance (HT). **a** Simulated mortalities (grey dots) of virtual corymbose acroporids assigned with a HT value (°C-week) generated at random from the initial distribution (μ=0, sd=2.49) and for random cumulative heat stress values (DHW). The blue and orange DHW/mortality curves indicate, respectively, the bleaching response of the 10^th^ and 90^th^ percentiles of this distribution, which reproduce the difference of 4.84°C-week inferred at the cut-off mortality of 12% (red dots) among the experimental samples of *Acropora digitifera* under heat stress^23^. The maximum difference (6.37°C-week) is achieved at 100% mortality (black dots). **b** Normal distribution of HT values of corals within each of the 6 modelled groups at initialisation (grey curve) with one possible realisation (grey histogram bars). Dotted lines indicate key percentiles, with the 10^th^ and 90^th^ corresponding to the experimental distribution of HT values extrapolated at 100% mortality. Red dotted lines indicate possible hard evolutionary limits of heat tolerance (±HTmax = 8°C-week), i.e., the lower and higher HT values that can be generated when a new coral is created. A coral with a HT value of 0°C-week (mean of the HT distribution, black line) has a mortality that follows the mean bleaching response of the group at initialisation (black curve in **a**).

**Extended Data Fig. 4.**
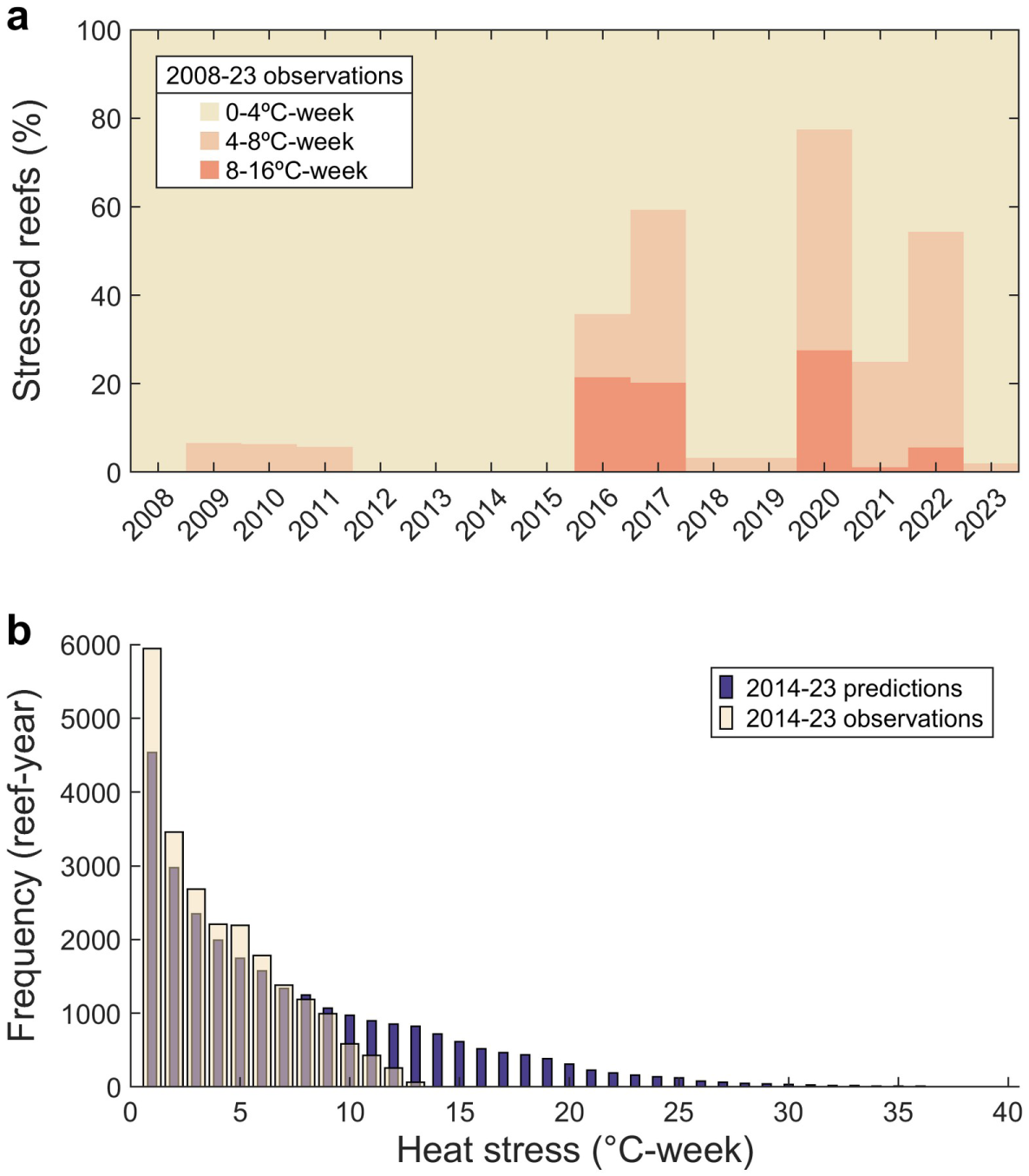
Recent heat stress on the Great Barrier Reef. **a** Proportion of individual reefs (n=3,806) under different categories of heat stress (DHW; °C-week) from 2008 to 2023 derived from 5-km resolution satellite SST observations (NOAA Coral Reef Watch^22^). **b**, Comparison of satellite observations vs. CMIP6 model predictions of reef-level DHW between 2014-2023 (n = 3,806 reefs × 10 years). Predicted (i.e., hindcast) DHW frequencies were averaged across the ten CMIP6 climate models and the five SSPs using the Equilibrium Climate Sensitivities as weights (see Methods).

**Extended Data Fig. 5.**
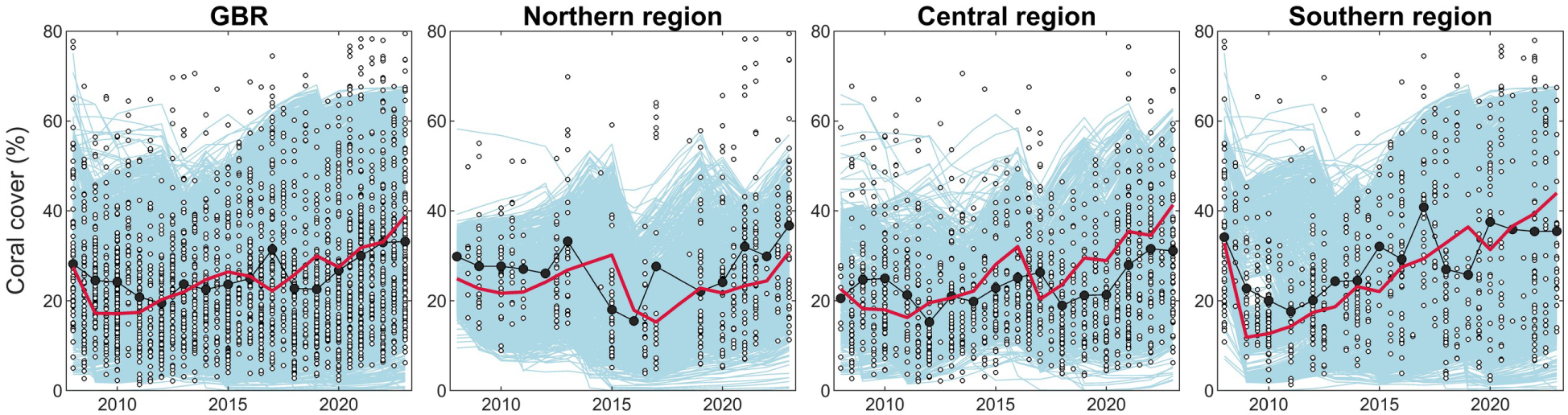
Hindcast (2008-2023) reconstruction of trajectories of total coral cover (blue lines: individual reef trajectories averaged over 20 simulations; red line: regional average) for the entire Great Barrier Reef (GBR, n = 3,806 reefs) and for the northern (n = 1,201 reefs), central (n = 957 reefs) and southern (n = 1,648 reefs) regions. Data points represent coral coverage observations from the Long-Term Monitoring Program of the Australian Institute of Marine Science^29^ (photo-transect and transformed manta tow estimates following ref. ^21^), where open dots correspond to individual reef surveys conducted during a specific season and filled dots indicate the annual average across all surveys. Average coral cover was not calculated for years with fewer than five reef surveys. An annual model error was calculated as the difference between the predicted and observed mean coral cover for each available year between 2009 and 2023 (excluding the 2008 observations that were used to initialise the simulated reefs), resulting in a mean (±SD) annual error of –0.3% (± 4.7%), –3.1% (± 8.6%), +1.7% (± 5.4%) and –3.3% (± 5.6%) for the GBR-wide, northern, central and southern reconstructions, respectively.

**Extended Data Fig. 6.**
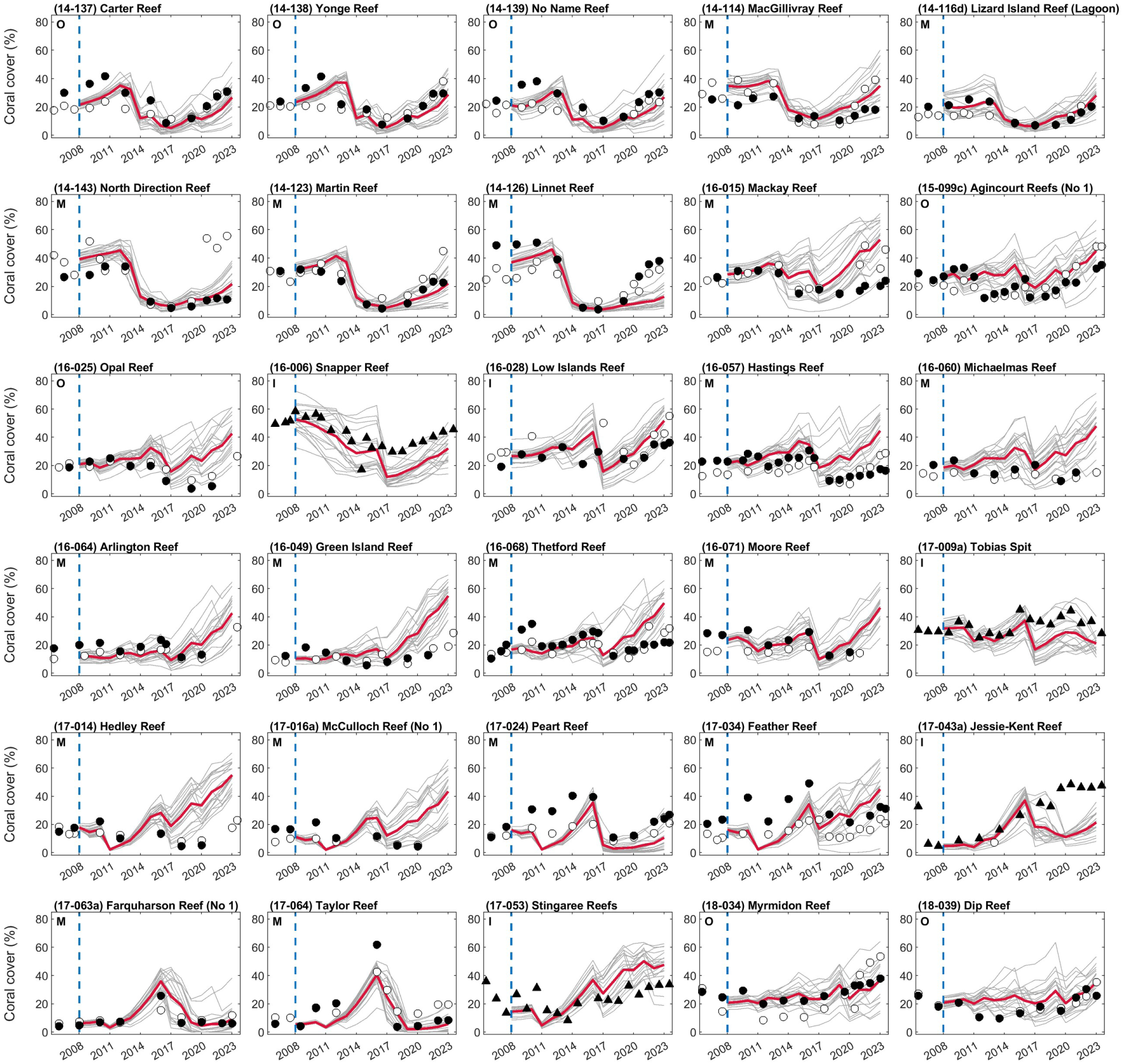
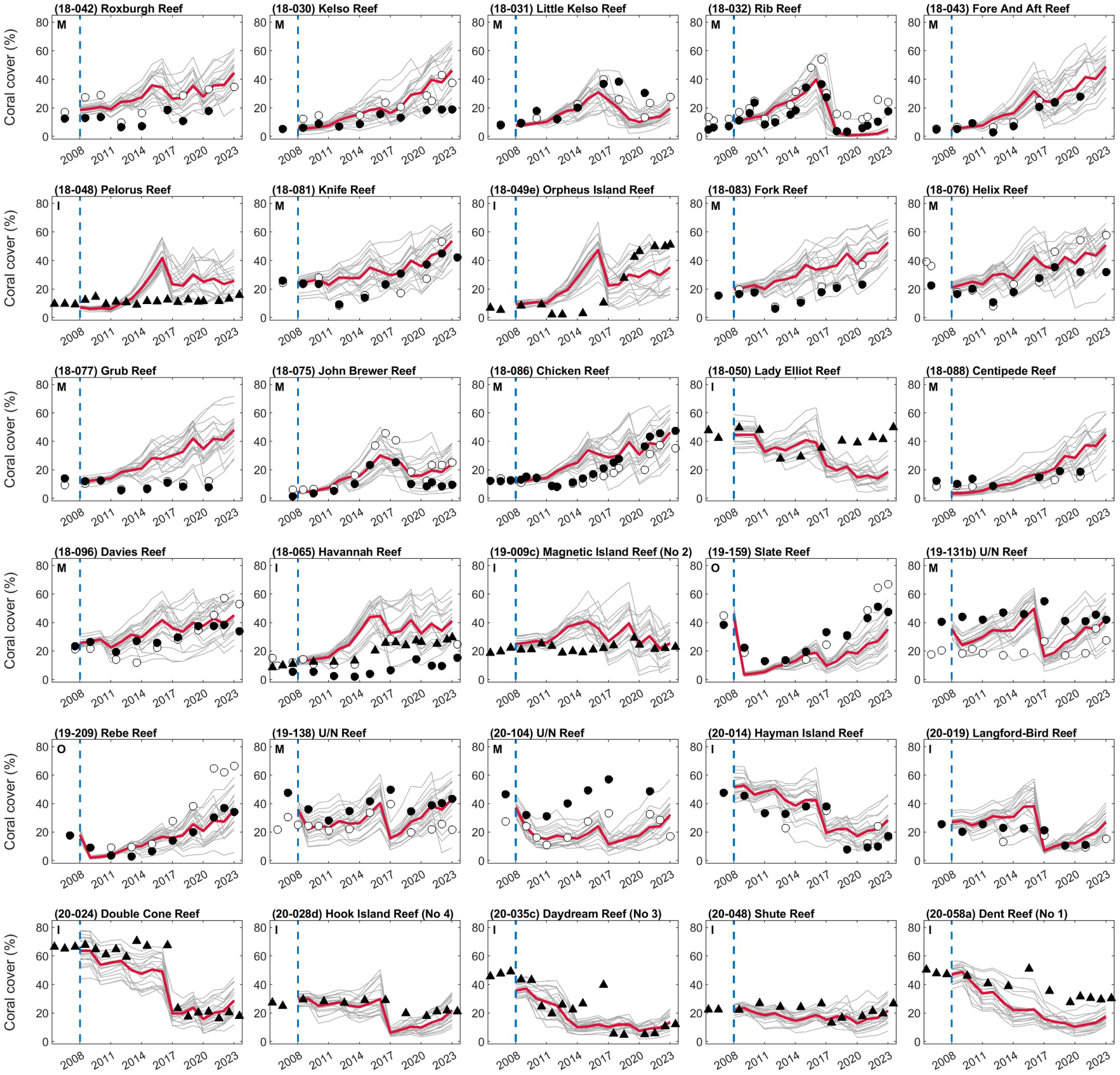
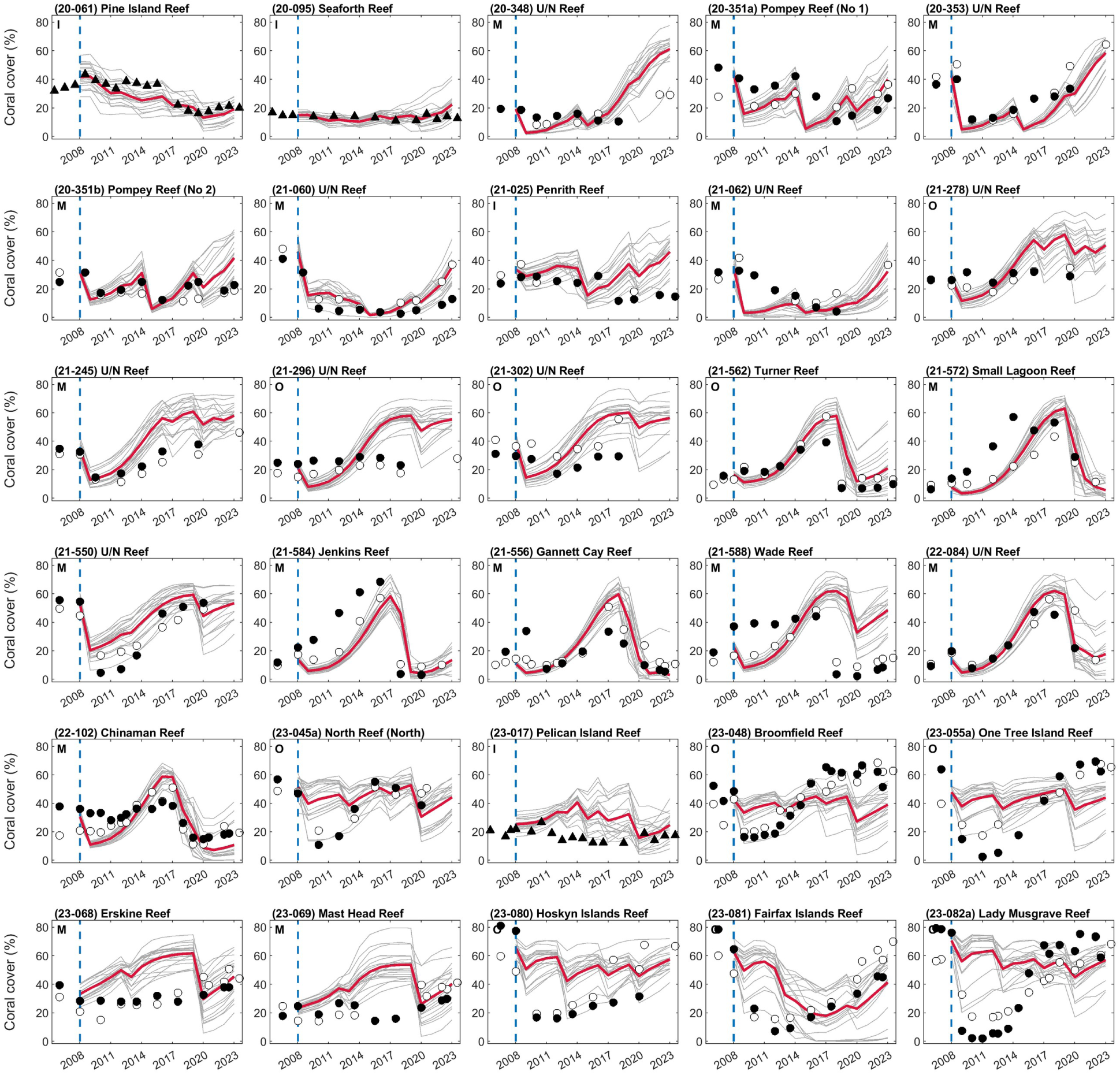
Hindcast (2008-2023) reconstruction of coral cover trajectories (grey line: trajectory of one stochastic simulation, N=20; red line: mean trajectory over all simulations) at individual reefs targeted by the Long-Term Monitoring Program (LTMP) and Marine Monitoring Program (MMP) of the Australian Institute of Marine Science^29^ (black dots: LTMP fixed photo-transects; open dots: LTMP manta tows, corrected to transect-equivalent coral cover, see ref. ^21^; black triangles: MMP fixed photo transects at 5 m depth). The selected reefs had at least one survey between 2006-2008, which is used to initialise coral cover in 2008 (blue dotted line), followed by at least nine surveys after 2008, to ensure a sufficient number of observations. Reefs are ordered from top to bottom according to their latitude (indicated by the first two numbers of their code label), with northernmost reefs at the top and southernmost reefs at the bottom. I: inshore; M: mid-shelf; O: outer-shelf.

**Extended Data Fig. 7.**
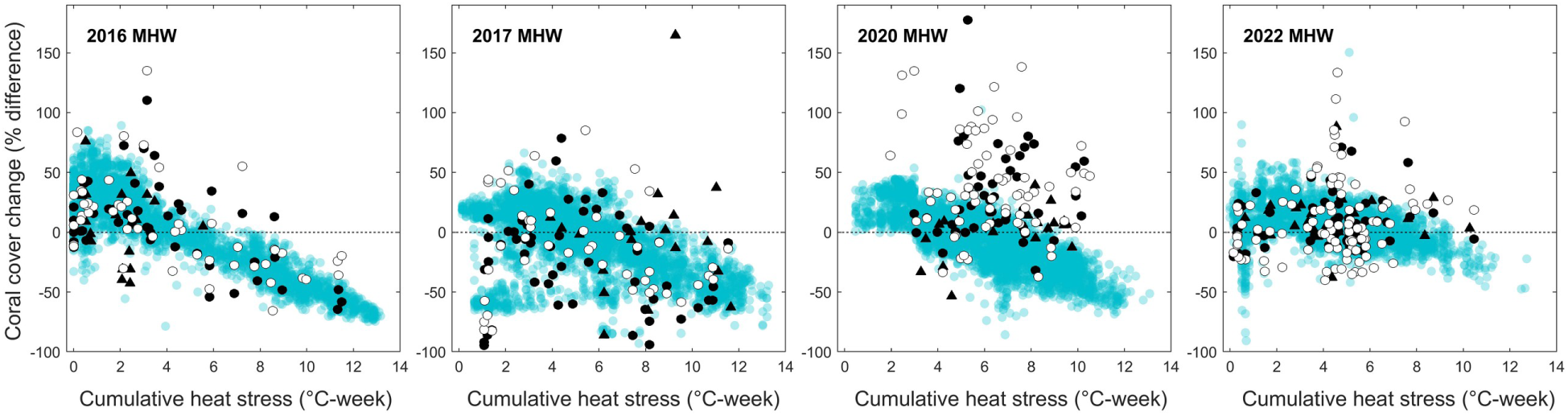
Reef-level relative changes in total coral cover following marine heatwaves (MHW) that occurred in 2016 (same as Fig. 1d), 2017, 2020 and 2022 across the GBR (blue dots, N=3,806 reefs) versus DHW exposure derived from satellite observations (NOAA Coral Reef Watch^22^). Coral cover changes were captured over a one-year interval. Positive coral cover changes likely indicate the absence of disturbance during the time period, resulting in growth and recruitment processes outweighing natural mortality. Negative coral cover changes underscore the impact of any disturbance, not just marine heatwaves. Note that bleaching was not simulated when reefs were exposed to 0-3°C-week heat stress^9^. The relative coral cover changes observed on reefs targeted by the Long-Term Monitoring Program (LTMP) and Marine Monitoring Program (MMP) of the Australian Institute of Marine Science^29^ are shown for comparison. Black dots: LTMP fixed photo transects; open dots: LTMP manta tows (corrected to transect-equivalent coral cover, see ref. ^21^); black triangles: MMP fixed photo transects (5 m depth). The dataset includes reefs that were monitored within one year before the MHW and again within one year after, allowing us to retain a significant amount of observations (2016 MHW: N=117 reefs; 2017 MHW: N=119 reefs; 2020 MHW: N=138 reefs; 2022 MHW: N=192 reefs).

**Extended Data Fig. 8.**
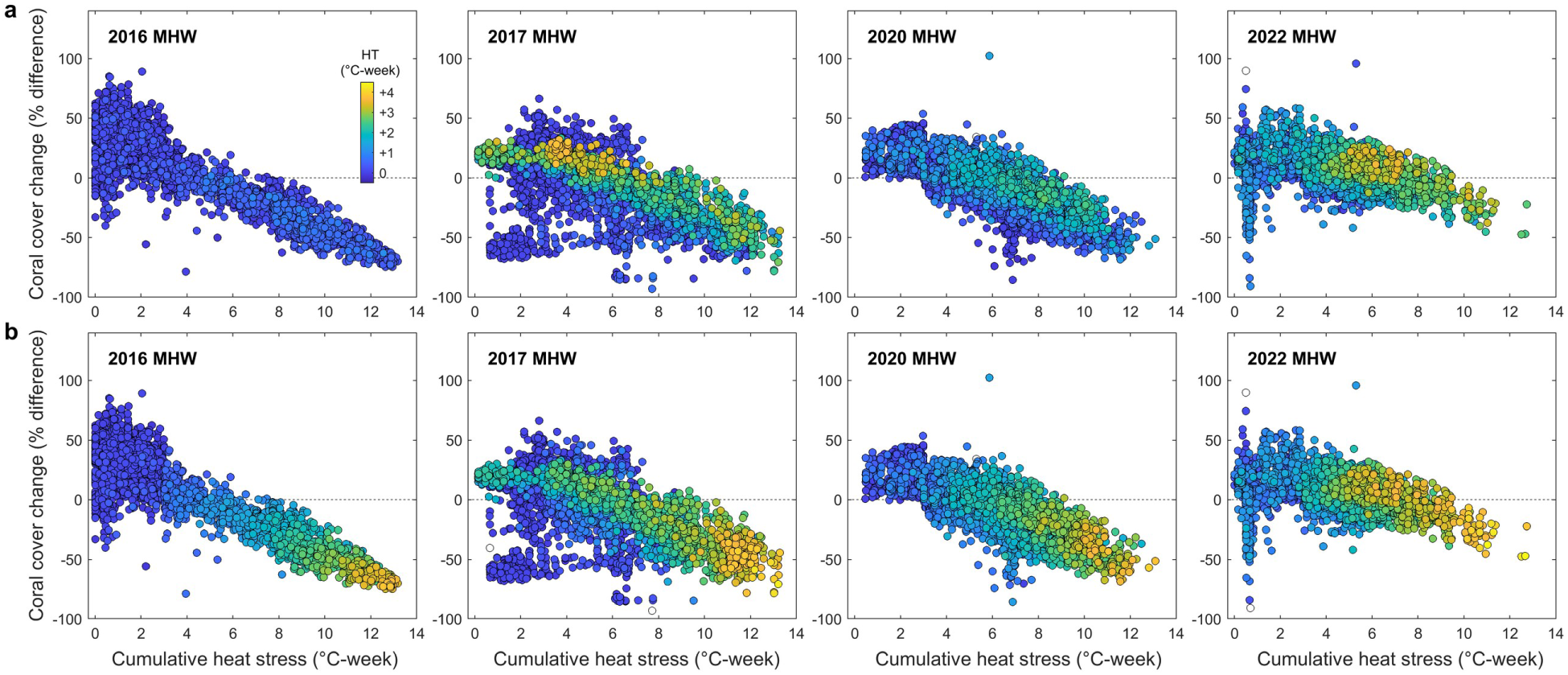
Reef-level relative changes in total coral cover following the 2016, 2017, 2020 and 2022 marine heatwaves as in Extended Data Fig. 7, showing changes in the mean heat tolerance (HT) of the most sensitive coral groups (acroporids and pocilloporids) before (**a**) and after (**b**) exposure to heat stress. Empty dots indicate reefs lacking sexually-mature colonies in the selected coral groups, which prevents calculation of mean thermal tolerance.

**Extended Data Fig. 9.**
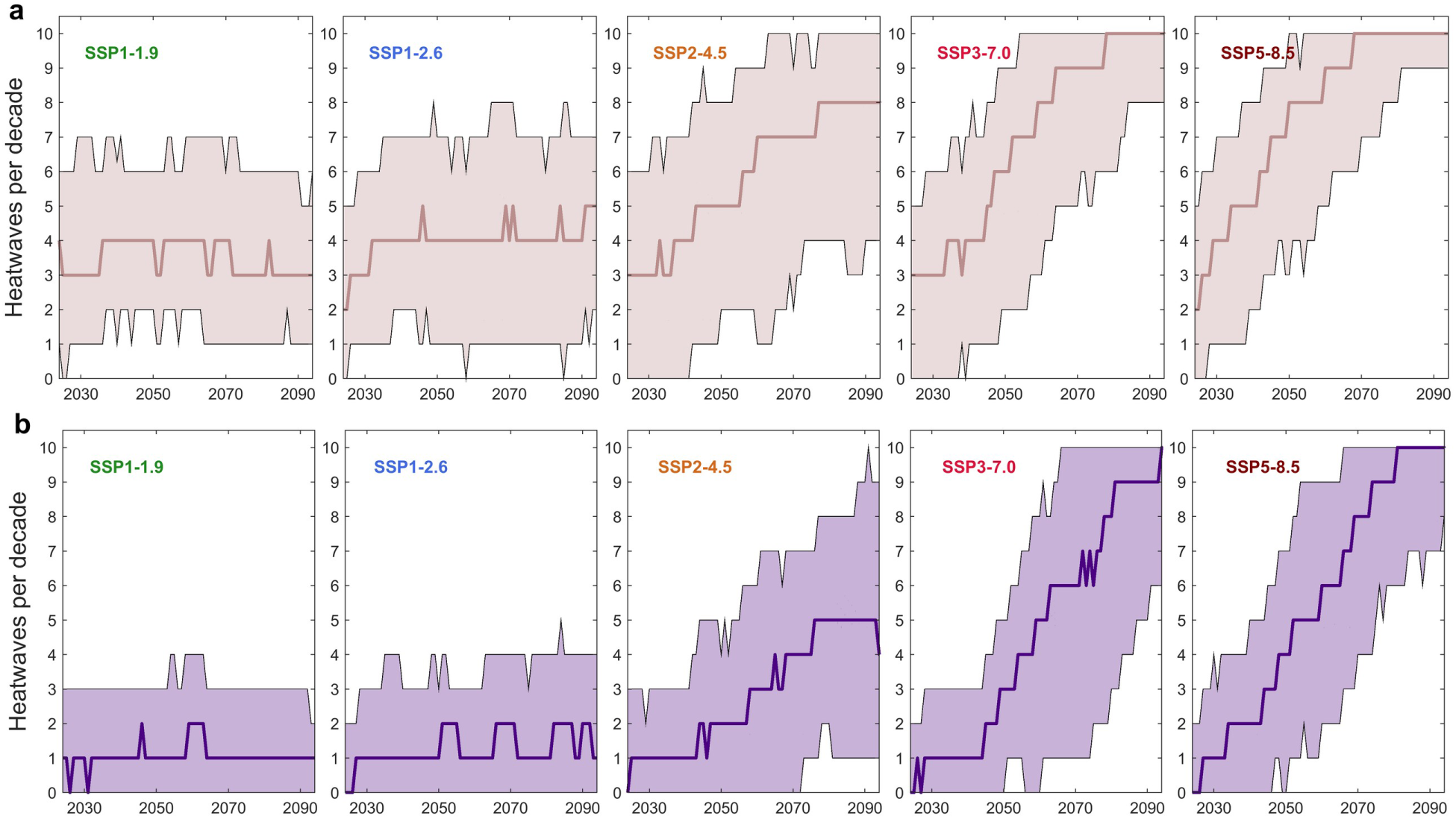
Reef-level decadal frequency of heat stress of severe intensity (**a**, DHW > 8°C-week; **b**, DHW > 16°C-week) experienced across the Great Barrier Reef. Thick lines indicate the median number of heat stress events above the two thresholds, while the lower and upper boundaries of the envelop represent the 10^th^ and 90^th^ percentiles, respectively (n = 3,806 reefs × 200 warming futures for each SSP, except for SSP1-1.9). The number of heat stress events above the specified threshold was calculated over a 10-year sliding window for every reef.

**Extended Data Fig. 10.**
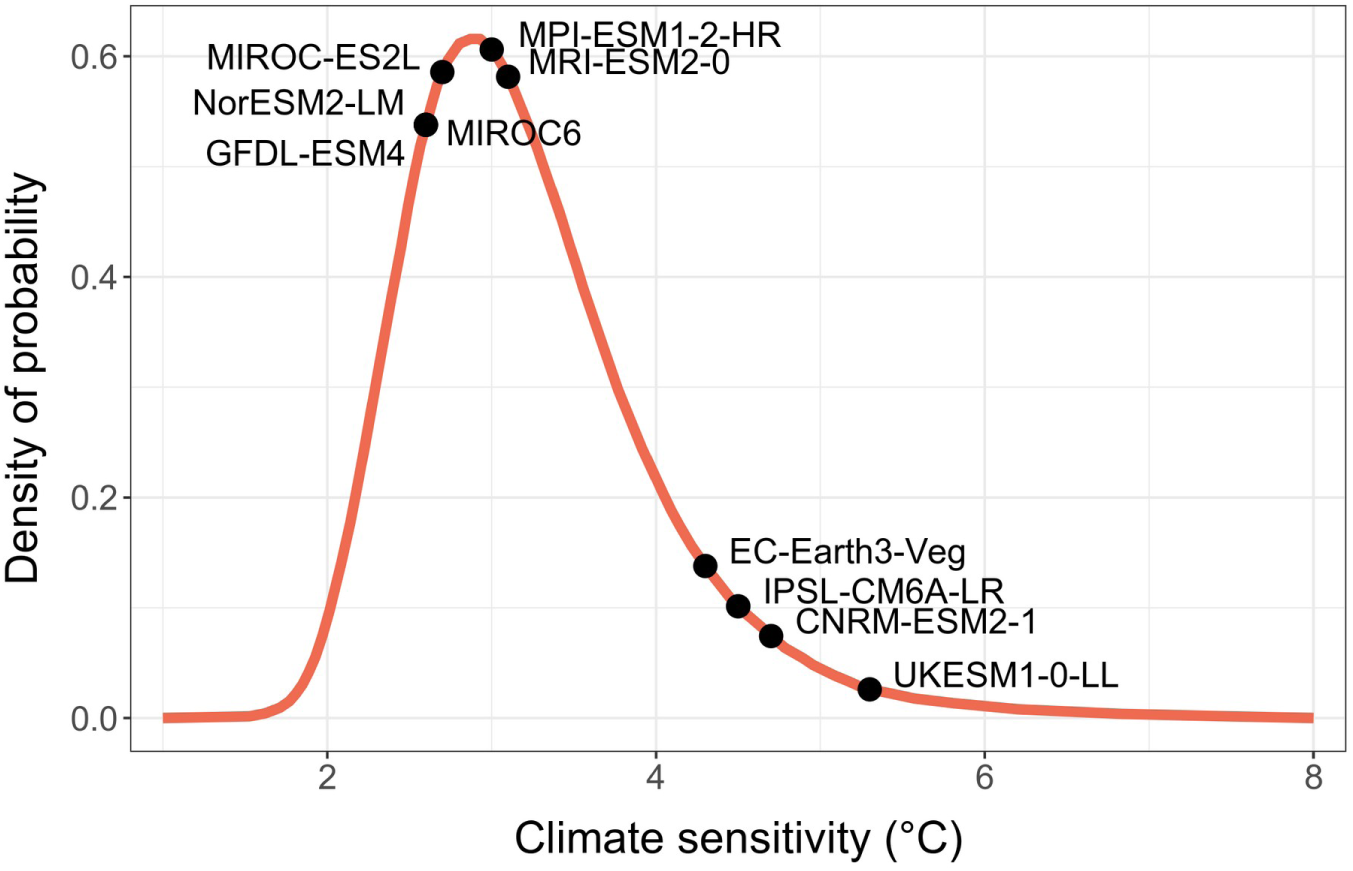
Probability density of Earth’s climate sensitivity (extracted from ref. ^86^) and equilibrium climate sensitivity (ECS) values of the ten CMIP6 climate models^38,84^.

**Extended Data Fig. 11.**
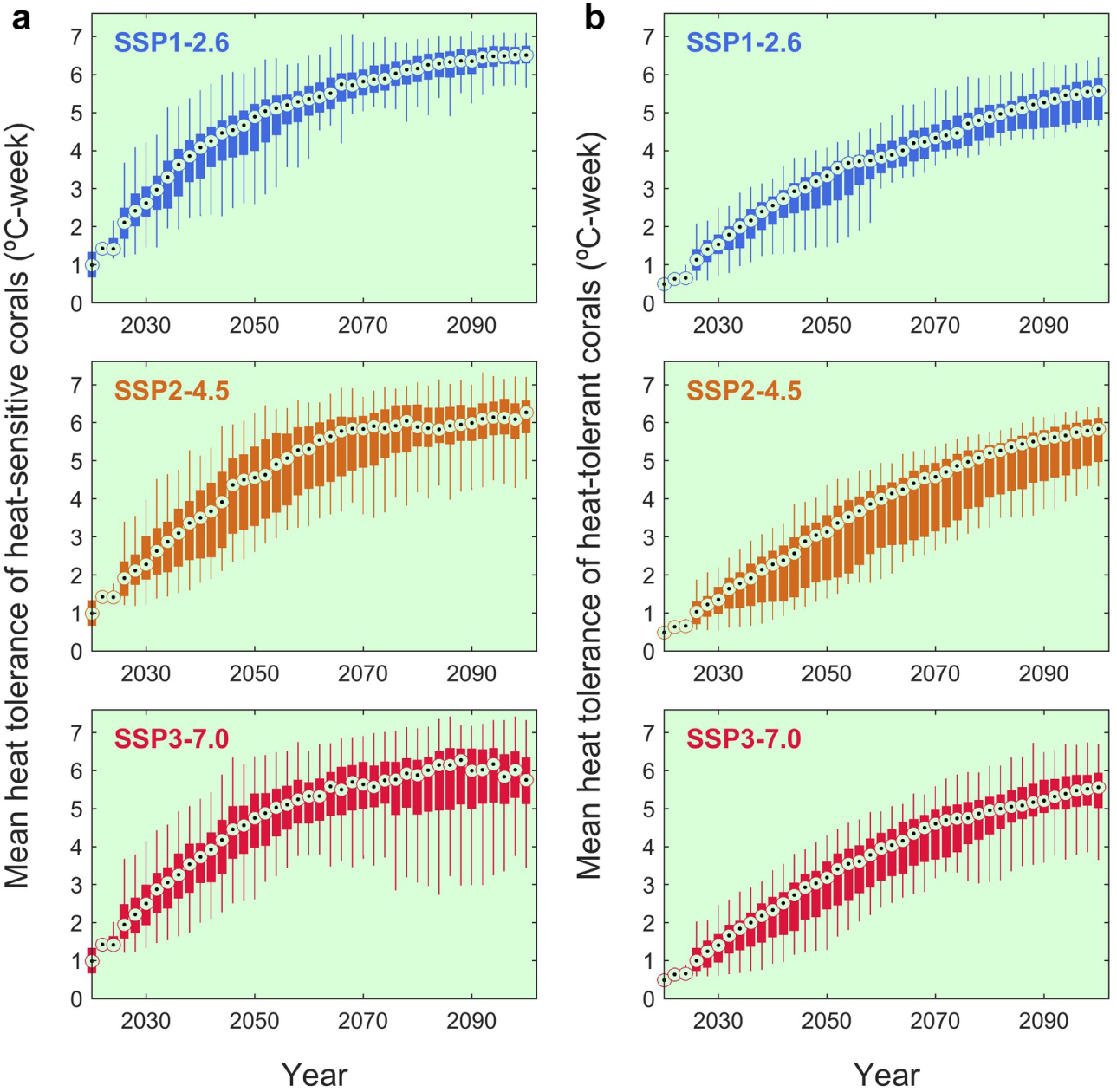
Great Barrier Reef mean heat tolerance of corals under the three most likely warming futures. From top to bottom, predictions are shown under SSP1-2.6, SSP2-4.5, SSP3-7.0 for (**a**) heat sensitive and (**b**) heat tolerant corals. Heat tolerance is defined as the relative accumulated heat stress (± °C-week) a coral colony can withstand compared to the mean response of its taxonomic group defined at initialisation. See Fig. 2 legend for details on the distributions.

**Extended Data Fig. 12.**
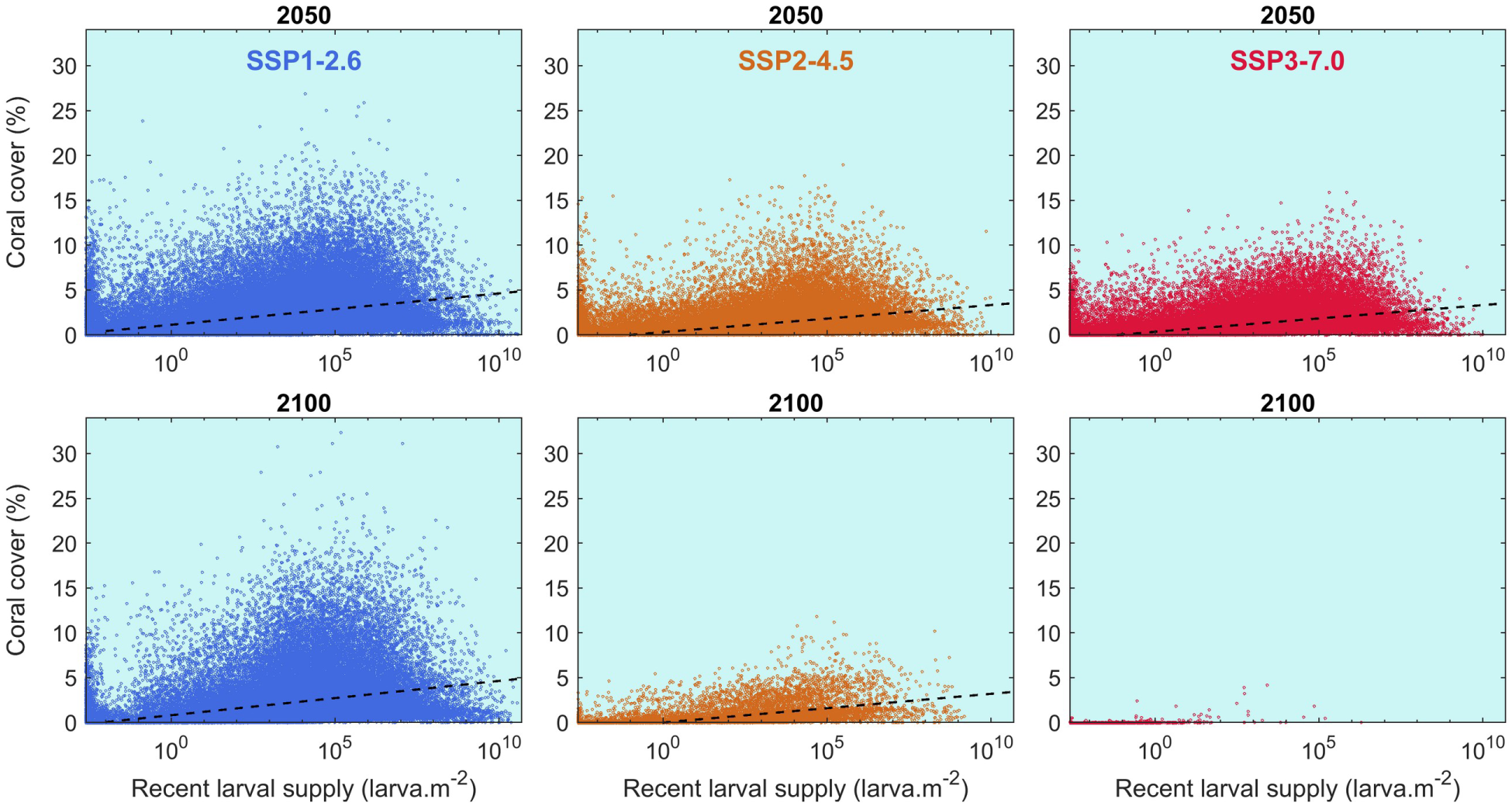
Influence of external larval supply on coral health in warm spots. The influence of external larval supply on coral cover closely depends on the context of recent exposure to disturbances, which can generate a diverse range of coral states making it difficult to depict a relationship between larval supply and coral cover. Focusing on warm spots (defined as the 90th percentile for mean annual DHW of the preceding 5 years) enlightens the relationship between coral cover on a reef and the average amount of external larvae supplied to that reef in the preceding 5 years.

**Extended Data Fig. 13.**
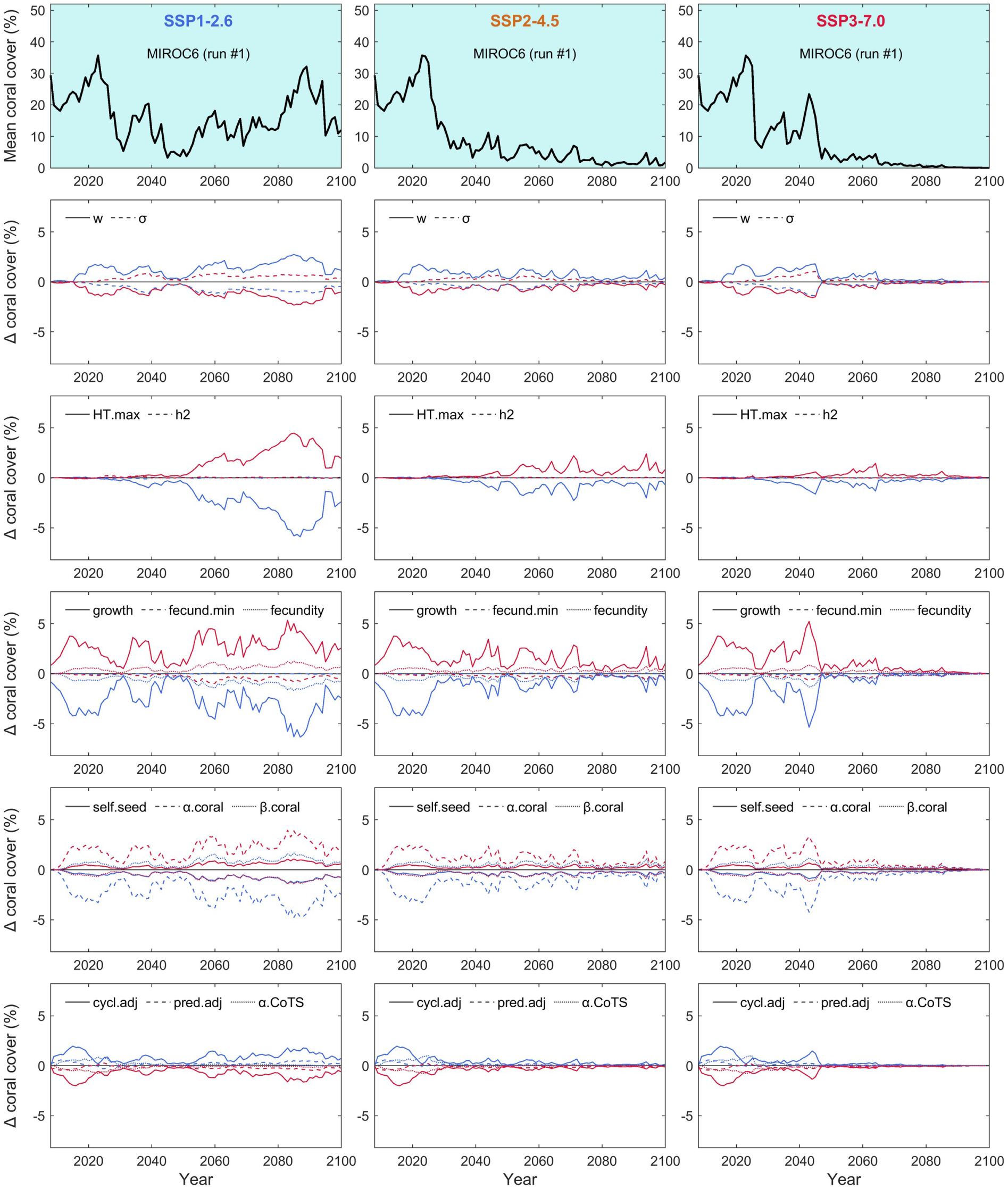
Sensitivity analysis. Model sensitivity to key input parameters (Extended Data Table 2) measured as the difference in the GBR mean coral cover between a simulation where one parameter is varied by ± 20% from its default value (blue: –20%; red: +20%) and the baseline simulation (top row) with all parameters at their default values. The baseline represents one scenario of future cyclones (run #1) across three SSPs (from left to right) as projected by the MIROC-6 climate model.

**Extended Data Table 1.** Predictors of reef state in mid and late century. State is defined as total coral cover across six diverse coral types. All metrics referenced to either the year 2050 (mid-century) or 2100 (late century).

| Predictor (as shown in figure) | Description | Unit | Time frame (years) |
| --- | --- | --- | --- |
| Years since bleaching | Number of years since last bleaching event | Years | Variable before each census date |
| Number of bleaching events | Total number of bleaching events | Events | 2025-2050 or 2075-2100 |
| DHW | Mean thermal stress where stress occurs when DHW≥6 | °C-week | 2025-2050 or 2075-2100 |
| Number of cyclone events | Total number of cyclone events | Events | 2025-2050 or 2075-2100 |
| Larval sink strength | Cumulative probability of larvae arriving at reef from other sources | 0-1 | Annual probability |
| Years since cyclone | Number of years since last cyclone event | Years | Variable before each census date |
| Cyclone strength | Mean category of all cyclones affecting the reef during census period | Cyclone category | 2025-2050 or 2075-2100 |
| CoTS density | Mean density of adult crown-of-thorns starfish per reef patch | Count per 400 m<sup>2</sup> | 2025-2050 or 2075-2100 |
| Number of CoTS control events | Number of control events in sampling period | Events | 2025-2050 or 2075-2100 |
| WQ impact on recruitment | Proportional coral recruitment after water quality effects where 0 is no recruitment and 1 recruitment unaffected | 0-1 | Fixed property of reef based on eReefs analysis (see ref. <sup>21</sup>) |
| Years since CoTS control | Number of years since last CoTS control | Years | Variable before each census date |
| WQ impact on reproduction | Proportional coral reproduction after water quality effects where 0 is no reproduction and 1 implies no impacts on reproduction | 0-1 | WQ impact on reproduction (see ref. <sup>21</sup>) |
| Larval supply in last 5 years | Cumulative larval supply in 5 years prior to census date, transformed by natural logarithm (+0.1) | Count | 2045-2050 or 2095-2100 |

**Extended Data Table 2.** List of parameters tested in the sensitivity analysis. Each parameter was varied by ± 20% of its default value (low/high figure), while keeping all other parameters at their default values.

| Parameter | Description | Default value | Low figure | High figure |
| --- | --- | --- | --- | --- |
| w | Depth-related attenuation coefficient of heat stress | 0.43 (7-m depth) | 0.34 (~10-m depth) | 0.52 (~5-m depth) |
| $\sigma$ | Standard deviation of heat tolerance within each coral group (naive distribution) | 2.5 °C-week | 2 °C-week | 3 °C-week |
| HTmax | Limits (negative and positive) to heat tolerance within each group | 8 °C-week | 6.4 °C-week | 9.6 °C-week |
| $h^2$ | Heritability (narrow-sense) of heat tolerance | 0.3 | 0.24 | 0.36 |
| Growth | Rate of lateral extension | 0.8 – 4.4 cm.yr<sup>-1</sup> | 0.6 – 3.5 cm.yr<sup>-1</sup> | 1.0 – 5.3 cm.yr<sup>-1</sup> |
| fecund_size | From Hall and Hughes 1996: size above which a colony is gravid<sup>89</sup> (taxon specific) | 31 – 134 cm<sup>2</sup> | 25 – 107 cm<sup>2</sup> | 37 – 161 cm<sup>2</sup> |
| fecund_b | Parameter *b* of the relationship linking colony size to total egg volume<sup>89</sup> (taxon specific) | 1.05 – 2.27 | 0.84 – 1.82 | 1.26 – 2.73 |
| Minimum larval retention | Minimum proportion of coral larvae produced on a reef that is retained for recruitment<sup>21</sup> | 0.28 | 0.22 | 0.34 |
| $\alpha$ | Maximum achievable density of coral settlers for a 100% free space<sup>21</sup> (across all taxa) | 15 settlers.m<sup>-2</sup> | 12 settlers.m<sup>-2</sup> | 18 settlers.m<sup>-2</sup> |
| $\beta$ | Density of coral larvae required to produce half the maximum settlement<sup>21</sup> (across all taxa). | 75,000 larvae.m<sup>-2</sup> | 60,000 larvae.m<sup>-2</sup> | 90,000 larvae.m<sup>-2</sup> |
| cycl_adjust | Global multiplier of cyclone sensitivity coefficients (obtained after calibration<sup>21</sup>) | 5 | 4 | 6 |
| pred_adjust | Global multiplier of CoTS individual consumption rate (obtained after calibration<sup>21</sup>) | 1.8 | 1.4 | 2.2 |
| $\alpha$ (CoTS) | Maximum achievable density of CoTS settlers (obtained after calibration<sup>21</sup>) | 100 settlers.m<sup>-2</sup> | 80 settlers.m<sup>-2</sup> | 120 settlers.m<sup>-2</sup> |

